# Internal friction can be measured with the Jarzynski equality

**DOI:** 10.1101/613091

**Authors:** R. Kailasham, Rajarshi Chakrabarti, J. Ravi Prakash

**Author notes:** Electronic mail (Rajarshi Chakrabarti).

## Abstract

A simple protocol for the extraction of the internal friction coefficient of polymers is presented. The proposed scheme necessitates repeatedly stretching the polymer molecule, and measuring the average work dissipated in the process by applying the Jarzynski equality. The internal friction coefficient is then estimated from the average dissipated work in the hypothetical limit of zero solvent viscosity. The validity of the protocol is established through Brownian dynamics simulations of a single-mode spring-dashpot model for a polymer. Well-established single-molecule manipulation techniques, such as optical tweezer-based pulling, can be used to implement the suggested protocol experimentally.

## I. Introduction

Conformational transitions in polymer molecules are impeded by solvent molecules and by intramolecular interactions within the molecule.^1–5^ The resistive effect from various intramolecular interactions^6–14^ is collectively called “internal friction”. Internal friction modulates the kinetics of spatial reorganization in a number of biological contexts, such as damping the process of protein folding,^14–17^ and affecting stretching transitions in polysaccharides.^6^ A commonly accepted operational definition for internal friction^14,16,18^ is the folding (reconfiguration) time for a protein in the extrapolated limit of zero solvent viscosity (*η*_s_ → 0). However, such an indirect definition does not ascribe a friction coefficient to the phenomenon. On the other hand, stretch-relaxation experiments on condensed DNA by Murayama et al.,^7^ and molecular dynamics simulations of polypeptide-stretching by Schulz, Miettinen and Netz^9^ provide an estimate for the internal friction coefficient. We are motivated by the protocol proposed by Netz and coworkers,^8,9^ wherein the work done in stretching a polymer molecule is partitioned into a reversible component, and an irreversible, dissipative component. The former goes into reversibly increasing the free energy, and is stored in the extension of the molecule. The latter represents the work expended in overcoming the rate-dependent restoring force offered by the the solvent degrees of freedom, and sources of friction internal to the molecule. In the model system examined in ref. 9, Netz and coworkers consider intramolecular hydrogen bonds as the source of internal friction. In the hypothetical limit of zero solvent viscosity, the only source of dissipation in the system is assumed to be the internal friction in the molecule. The average dissipated work in the limit of zero solvent viscosity, is then used to estimate the internal friction coefficient, and is found to scale with the number of intramolecular hydrogen bonds in the system. In this paper, we propose a methodology by which single molecule stretching experiments using optical tweezers could be employed in conjunction with a novel application of the Jarzynski equality^19^ to measure the dissipation associated with the stretching process and consequently extract the internal friction coefficient of the molecule. The Jarzynski equality provides a conceptual framework for obtaining the free-energy difference associated with a process from a non-equilibrium protocol. Since its formulation in 1997, the Jarzynski equality has been routinely employed for reconstructing the free energy landscape of biomolecules from finite-rate pulling experiments^20–22^ and simulations.^23,24^ The dissipation arising due to the non-equilibrium protocols employed in these applications has largely been ignored, except in the context of estimating the accuracy^25–27^ of the free-energy difference obtained using the Jarzynski equality. In this study, we leverage the dissipated work to our advantage, and show that internal friction can be measured with the Jarzynski equality.

It is worth noting that prior studies on the dissipative effects of internal friction^7–9^ calculate the reversible and irreversible components of the total stretching work *separately*: the free energy difference is obtained as the work done in the quasi-static limit,^28^ and the dissipated work at a given pulling rate is obtained by subtracting the free energy difference from the total work done. Here we propose that multiple realizations of the pulling experiment be performed, which yield a distribution of work values due to thermal fluctuations in the system. From this distribution of work values, the Jarzynski equality allows one to extract the free-energy difference and the average dissipated work *simultaneously*, at *finite* pulling rates. At a fixed value of the pulling velocity, the average dissipated work is then calculated at various values of the solvent viscosity, *η*_s_, and its value in the hypothetical limit of zero solvent viscosity, 〈*W*_dis_〉_*η*_s_ → 0_, is obtained by extrapolation. This exercise is to be repeated for multiple values of the pulling velocity. The dissipative force due to internal friction is given by the ratio of 〈*W*_dis_〉_*η*_s_ → 0_ to the distance, *d*, over which the molecule is stretched. The internal friction coefficient is then evaluated as the slope of the plot between the dissipative force and the pulling velocity.

The validity of this protocol is established through Brownian dynamics (BD) simulations. A spring-dashpot model for a polymer is considered, where the molecule is represented as two massless beads connected by a spring in parallel with a dashpot (see Fig. 1 (a)), subjected to fluctuating random forces from the bath of solvent molecules it is immersed in. This model has been widely invoked in rheological,^1,29–31^ as well as biophysical contexts,^6,12,32^ to capture the effects of internal friction in polymers. Within this model, the spring accounts for the entropic elasticity in the polymer molecule, whereas the dissipative effect from the myriad sources of internal friction is captured by the dashpot. The solvent drag on the beads accounts for the friction caused by the motion of the polymer in the solvent. The solvent-mediated propagation of momentum between the beads is accounted for by the inclusion of fluctuating hydrodynamic interactions (HI), since it is well established that fluctuations in HI affect the dynamic response of polymer molecules.^31,33^ The present study is the first derivation and numerical solution of the system of stochastic differential equations necessary for carrying out force spectroscopy simulations of the spring-dashpot model. Using this model, we demonstrate that the internal friction coefficient estimated from the average dissipated work in the hypothetical limit of zero solvent viscosity is identical to the damping coefficient of the dashpot, which is a model input parameter. The successful recovery of the damping coefficient suggests that the proposed protocol could be used to experimentally obtain an estimate of the magnitude of the internal friction coefficient in polymers.

**FIG. 1.**
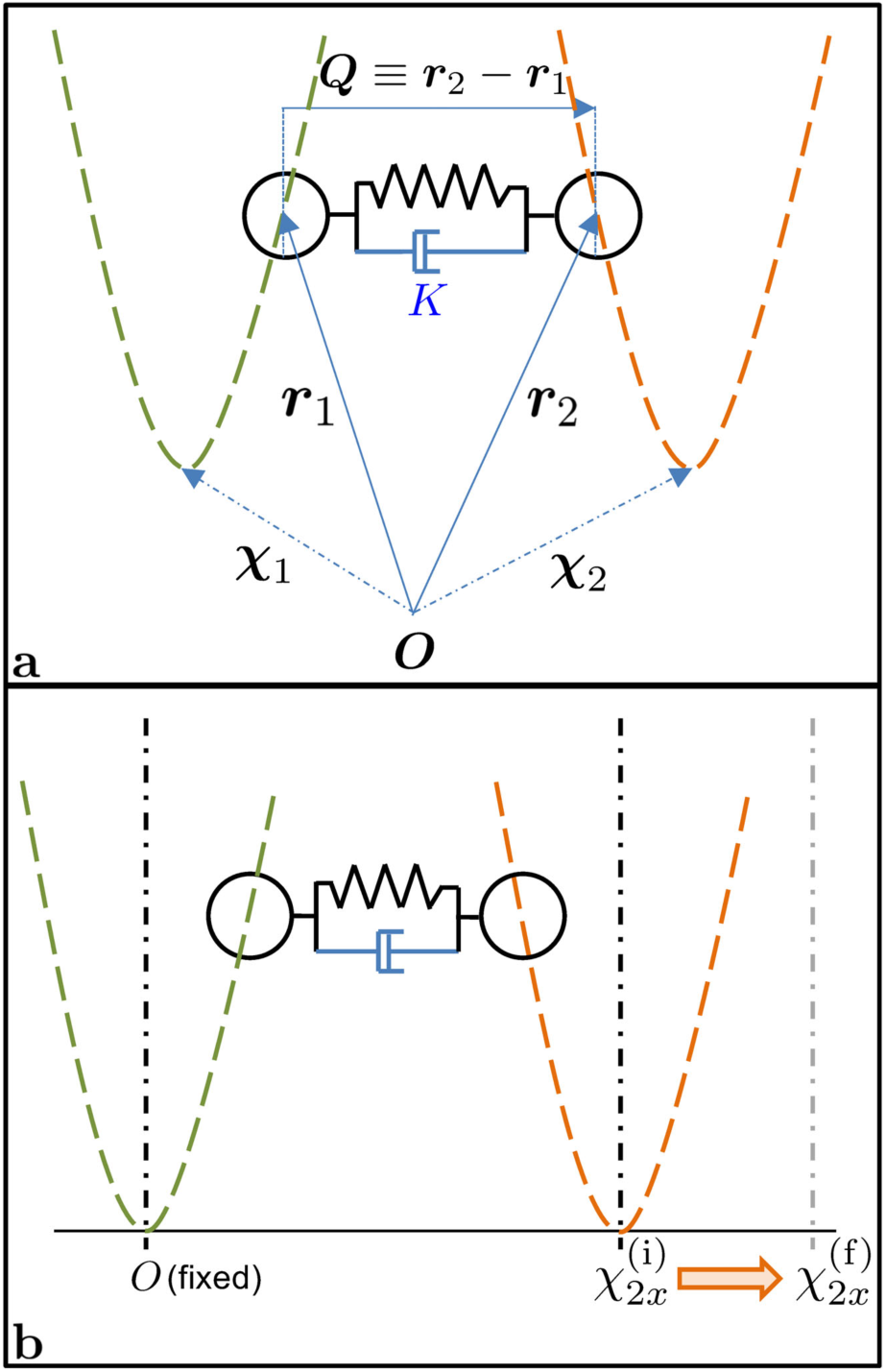
Simulation schematic/Proposed experiment. (a) Schematic diagram of the coarse-grained polymer model entrapped between two optical tweezers. The spring connecting the two beads is finitely extensible, upto a length *Q*_0_. Internal friction is modelled using the dashpot, whose damping coefficient is *K*. The Hookean spring constant associated with the spring is *H*. The strengths of the two traps, modelled as harmonic potential wells, are *H*_1_ = *c*_1_*H* and *H*_2_ = *c*_2_*H* respectively. (b) The one-dimensional pulling protocol: the position of the first trap is taken to be the origin, and remains stationary throughout the experiment. The second trap is moved from its initial position, 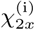 to its final position, 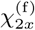, over a time-interval *τ*, stretching the spring-dashpot setup in the process. The difference between the initial and final positions of the mobile trap is *d*, and the velocity of pulling is *v*_*x*_.

While the proposed ansatz rests on an assumption that awaits rigorous proof, namely, that the total friction experienced by the polymer molecule may be separable into a “dry” component (independent of solvent viscosity) and a “wet” component (arising from solvent drag), a recent study by Daldrop et al.^34^ in which they calculate the total friction using a generalized Langevin equation that incorporates both inertial and solvent memory effects, lends substantial credence to this empiricism. In the present treatment, an overdamped Langevin equation is considered that ignores both inertial and memory effects. Furthermore, it is believed that force-induced unfolding in general follows a different mechanistic pathway in comparison to those induced by chemical agents^35^ or temperature.^36^ Consequently, the internal friction coefficient determined from stretching experiments might not necessarily represent the friction associated with the global unfolding of the molecule initiated by chemical or thermal means. Nevertheless, in contexts where it is applicable, the proposed protocol offers a means of quantifying the magnitude of internal friction directly, unlike the majority of the approaches used so far.

The rest of the paper is organized as follows. After introducing the model and the pulling scheme used in our simulations, a brief description of the BD simulation procedure is provided. The parameters used in the simulations, and the rationale behind their choice is then explained. The protocol for the extraction of the internal friction coefficient, and the results are then presented. Finally, we offer a few concluding remarks.

## II. Model description

As shown in Fig. 1 (a), we consider a dumbbell model for a polymer with fluctuating internal friction and hydrodynamic interactions. The beads (each of radius *a*) are joined by a spring, with maximum stretchability *Q*_0_ and a Hookean spring constant *H*, in parallel with a dashpot of damping coefficient *K*. The dumbbell is suspended in an incompressible, Newtonian solvent of viscosity *η*_s_. The Marko-Siggia force expression,^37^ widely employed to model the force-extension relationship in synthetic polymer molecules,^38^ as well as biopolymers,^39–42^ is used to describe the entropic elasticity in the dumbbell. The connector vector joining the two beads is denoted by ***Q*** ≡ ***r***_2_ − ***r***_1_. The centre-of-mass coordinates are given by ***R*** ≡ (1/2) (***r***_1_ + ***r***_2_). The positions of the two beads can be manipulated using optical traps, modelled here as harmonic potential wells. The trap stiffnesses are denoted by *H*_1_ = *c*_1_*H*, and *H*_2_ = *c*_2_*H* (in units of the dumbbell spring constant), and the co-ordinates of the minimum of the wells are represented by ***χ***_1_ and ***χ***_2_, respectively. A temperature of *T* = 300 *K* is considered in all our simulations, as a matter of convenience. The viscosity of the solvent at this temperature is taken to be *η*_s,0_ = 0.001 kg/m s, which is close to the viscosity of water at room temperature. In this protocol, values of solvent viscosity which are multiples of *η*_s,0_ will be considered. Within this model, the conservative spring potential acting between the beads of the polymer determines the equilibrium properties of the molecule. The dashpot does not affect equilibrium properties, and only modulates the dynamics of the polymer molecule.^31^ A dashpot of damping coefficient *K* pulled with a constant velocity *v* over a distance *d*, offers a resistive force of magnitude *Kv*.^1,43^ The work done against this restoring force is then simply given by the product of the restoring force and the stretching distance,

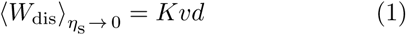

The temperature and the spring stiffness together define the length scale of the problem, 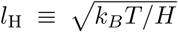, where *k*_*B*_ is the Boltzmann constant. Using this definition of the length-scale, a finite extensibility parameter, *b*, is introduced as 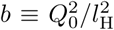. The molecular time scale is given by λ_H_ = *ζ*/4*H*, where *ζ*(:= 6π*η*_s_*a*) is the bead friction coefficient. The internal friction parameter, ϵ(:= 2*K*/*ζ*), is defined as the ratio of the internal friction coefficient to the bead friction coefficient. Quantities with an asterisk as superscript are understood to be dimensionless.

In Fig. 1 (b), the pulling protocol employed in this study is depicted. Without any loss of generality, ***χ***_1_ is chosen as the origin of our frame of reference. In all pulling simulations throughout this work, the first trap is held stationary, and the second trap is moved from its initial position, 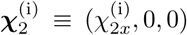, to its final position, 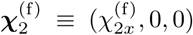. The notation 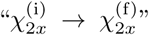 represents such a pulling event. The stretching distance is denoted by 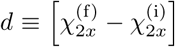, the time interval for stretching by by *τ*, and the pulling velocity by ***v*** ≡(*v*_*x*_, 0, 0), where *v*_*x*_ = *d*/*τ*. Note that the bead co-ordinates are allowed to sample the entirety of the three-dimensional coordinate space, but the pulling is restricted to the *x*-axis alone. One could implement an alternative protocol such that the pulling direction is also in general three-dimensional space. However, such a change will not alter the analysis and arguments presented in this study.

For any value of the trap position, ***χ***_2_, the Hamiltonian, 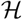, of the system is given by,

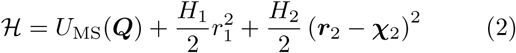

where *U*_MS_(***Q***) represents the potential energy in the spring. The generalized Jarzynski work^44^ corresponding to the pulling protocol discussed above is given by

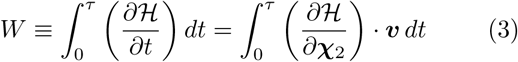

where ***v*** = *d****χ***_2_/*dt*.

The work done during each pulling event is different due to the thermal fluctuations in the system. The Jarzynski equality^19^ is used to evaluate the free energy difference, ∆*A*, corresponding to the transition shown in Fig. 1, as

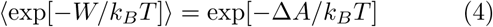

where 〈…〉 denotes an average over all possible realizations of the pulling protocol. The average work dissipated in the process is given by 〈*W*_dis_〉 = 〈*W*〉 − ∆*A*.

## III. Brownian dynamics simulations

We use Brownian dynamics to simulate the pulling depicted in Fig. 1 (see Sec. I of the SI for details of the governing equations and numerical scheme). Fluctuating hydrodynamic interactions are modeled using the Rotne-Prager-Yamakawa tensor.^45,46^ The hydrodynamic interaction parameter is given by 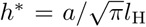, and *h\** = 0 corresponds to the free-draining case. The governing system of stochastic differential equations in ***Q^∗^*** and ***R^∗^*** is solved in its dimensionless form, using a semi-implicit predictor corrector method. The corresponding dimensional quantities are obtained by a suitable multiplication with the scaling factors.

A dimensionless time-step width of ∆*t^∗^* = 1 × 10^*−*3^ is used for all the parameter sets considered in this work. This value corresponds to the highest time-step width that gives converged results. Higher values of the internal friction parameter, smaller values of the finite extensibility parameter, or stiffer optical traps would necessitate the use of smaller ∆*t^∗^*, as elaborated in Sec. I of the SI.

The dimensionless equivalent of the work done during one realization of the pulling event is calculated by evaluating the integral in Eq. 3 using a simple rectangular quadrature (see Sec. I of the SI for details).

The free-energy difference between the initial and final states can be computed exactly, since the Hamiltonian of the system is known [Eq. 2]. There is no closed-form analytical solution for the partition function, as a consequence of the non-linearity introduced by the Marko-Siggia spring potential, and therefore, the free energy difference is evaluated using numerical quadrature. The free energy difference computed in this manner is compared with the result obtained from the implementation of Jarzynski’s equality using BD simulations, for two sample cases. Excellent agreement between the free energy differences computed using the two approaches establishes the validity of the code. Additional details pertaining to the validation studies are presented in Sec. II of the SI.

## IV. Parameter-space specification

The parameters used in the present work are broadly classified into molecular and control parameters. Molecular parameters pertain to the polymer that is being stretched, whereas control parameters are set by the experiments or the simulations used in the study of stretching the molecule.

### A. Molecular parameters

The molecular parameters relevant to this study are the finite extensibility parameter, *b*, the length scale, *l*_H_, the bead radius, *a*, and the damping coefficient of the dashpot, *K*. The dumbbell parameters are chosen such that they model the *λ*-phage DNA (contour length: 16.5*µ*m, Kuhn segment length: 88 nm) used by Murayama et al.^7^ in their stretch-relaxation experiments. As shown in Sec. 3A of the SI, the appropriate values in this case are *b* = 800 and *l*_H_ = 500 nm. Alexander-Katz et al.^8^ suggest that the monomeric radius may be taken as the persistence length of the molecule. A bead radius of *a* = 30 nm is chosen as a representative value. Additional values of *b*, *l*_H_, and *a*, of about the same order-of-magnitude, have also been used so as to test the protocol for a range of parameter values. The internal friction coefficient associated with the molecule used by Murayama et al.^7^ is *K* ≈ 10^*−*7^ kg/s. Molecular dynamics simulations of pulling experiments on polypeptides by Netz and coworkers^9^ estimate the internal friction coefficient to be in the order of ~ 10^*−*10^ kg/s. In this work, internal friction coefficients that fall within the range mentioned above are chosen, by picking values for *K* between 1.0 × 10^*−*9^ kg/s and 1.0 × 10^*−*8^ kg/s. Sec. 3A of the SI presents a detailed discussion on the molecular parameters selected for this study.

### B. Control parameters

The control parameters in the current study are: the stiffness of the optical traps, the initial extension of the molecule subjected to the stretching protocol, and the pulling velocity. The rationale behind the choice of these parameters is briefly discussed below.

#### 1. Trap stiffness

The strength of the optical trap determines its ability to confine the bead to positions close to its minimum. Higher trap stiffnesses ensure that the beads track the positions of the trap minima (Fig. S2 of the SI), but also necessitate the use of smaller time-step widths in our simulations. For a given set of molecular parameters, terminal states, pulling velocity, and stationary trap stiffness *c*_1_ = 1000, the dissipated work increases with an increase in the mobile trap stiffness, and saturates to a constant value at around *c*_2_ ≈ 100 (see Fig. S3 of the SI). Since it is desirable to operate in a regime where the dissipated work is independent of trap stiffness, *c*_1_ = *c*_2_ = 1000 is used in all our simulations.

#### 2. Initial stretch of molecule

The restoring force in the spring increases linearly at low values of the fractional extension 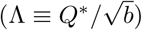, and diverges as Λ ~ 1 (Fig. S5 of the SI). As expected in a non-linear spring, stretching becomes increasingly harder at higher fractional extensions, necessitating the use of stiffer traps in order that the beads track the position of the traps. As shown in Sec. 3B of the SI, for a fixed value of the trap stiffness, increasing the ensemble size decreases the error in the recovered internal friction coefficient when the stretching is restricted to the linear regime. However, when the protocol is extended to the non-linear regime, higher values of the trap stiffness are required to recover the internal friction coefficient with the same accuracy. Throughout this paper, initial states of the spring such that Λ ≤ 0.2 have been chosen, but, as shown in Table S5 of the SI, the analysis is transferable to springs that start at Λ ~ 0.7, provided that stiffer traps are used.

#### 3. Pulling velocity and ensemble size

In Fig. 2, the average dissipated work (scaled by the thermal energy *k*_*B*_*T*) calculated for a variety of molecular and control parameters is plotted against the magnitude of the dimensionless pulling velocity, *v*^∗^. It is seen that the average dissipated work varies linearly over the entire range of the pulling velocity, *v*^∗^ = 0.001 − 0.1, except for the dataset with the highest value of the internal friction parameter, which deviates from linearity at higher pulling velocities. The velocity range in dimensional units would depend on the molecular parameters.

**FIG. 2.**
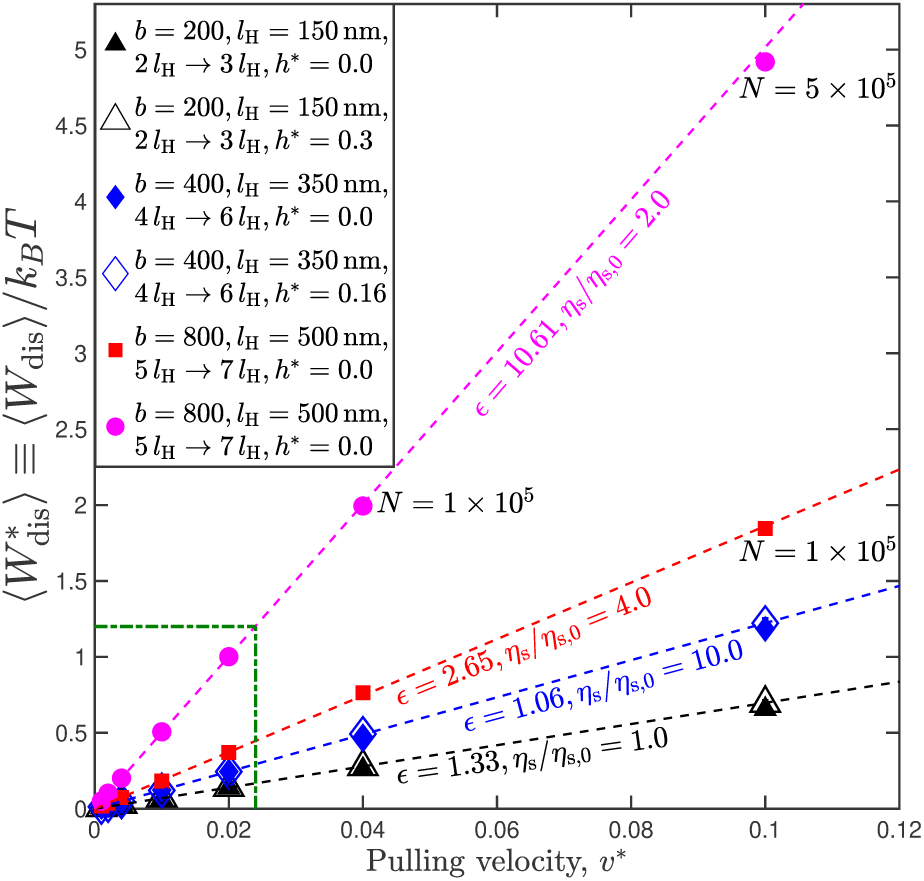
Linearity between dissipation and pulling velocity determines regime of operation. Average dissipated work as a function of the dimensionless pulling velocity, for various molecular and control parameters. Except when mentioned otherwise, an ensemble size of *N* = 1 × 10^4^ is used for all the data points. Symbols indicating datasets with fluctuating hydrodynamic interactions have been enlarged for the sake of clarity. Boxed region indicates the regime of operation throughout this paper. Error bars, which represent a statistical uncertainty of one standard error of the mean (s.e.m), are smaller than the symbol size.

The experimental feasibility of the proposed protocol can be discussed in the context of the molecular parameters used for the dataset represented by filled circles in Fig. 2. For these set of parameters, the pulling velocities explored in Fig. 2 vary from *v* = 29.3 nm/s (*v*^∗^ = 0.001) to *v* = 2.93 *µ*m/s (*v*^∗^ = 0.1). The molecule is stretched over a distance of 1*µ*m. The stiffness of this molecule is *H* = 1.657 × 10^*−*5^ pN/nm. In order to operate in a regime where the dissipated work is independent of the trap strength, as discussed earlier, the stiffness of the trap must be at least a hundred times that of the molecule, which implies *H*_trap,min_ = 1.657 × 10^*−*3^ pN/nm. These values of *v*, *d*, and *H*_trap,min_ lie well within the range of values explored experimentally, as discussed in greater detail in Sec. 3E of the SI.

Ritort et al.^25^ have established from computer simulations of mechanical unfolding that the number of trajectories required to obtain estimates for free energy difference within an error of 𝒪(*k*_*B*_*T*) increases exponentially with the average dissipation associated with the unfolding process. They predict that for dissipation less than 4*k*_*B*_*T*, around 100 trajectories would suffice, and for a dissipation of 5*k*_*B*_*T*, about 1000 trajectories would be required. These predictions agree well with the average dissipation and ensemble sizes encountered in optical-tweezer-based pulling experiments on RNA^20^ and DNA hairpins.^22^ In the present work, error in the free-energy difference is maintained to be ~ 𝒪 (0.01*k*_*B*_*T*), in order to obtain a sufficiently accurate estimate of the average dissipated work that enables the internal friction coefficient to be extracted reliably. By restricting the regime of operation to the boxed region in Fig. 2, with *v*^∗^ ≤ 0.02 and 〈*W*_dis_〉 ∼ *k*_*B*_*T*, it is found that *N* = 1 *×* 10^4^ trajectories are sufficient to ensure convergence in the free energy difference within the desired error-bounds. It is possible to operate at higher values of dissipation, outside the boxed regime, provided that the ensemble size is suitably increased. A detailed account on the choice of ensemble size is provided in Sec. 3D of the SI.

Netz and coworkers^8,9^ report a non-linear relationship between the dissipated work and the pulling velocity at high values of the latter quantity, and restrict their analyses to the linear regime for the extraction of the internal friction coefficient. Along similar lines, the current protocol relies on the linear relationship between dissipated work and pulling velocity in order to meaningfully define the internal friction coefficient.

Speck and coworkers^47,48^ have shown in the context of colloidal particles and a dumbbell model for a polymer, that the inclusion of hydrodynamic interactions does not alter the dissipation along a single trajectory. It is observed here as well, that the inclusion of fluctuating hydrodynamic interactions does not affect the dissipated work values, as seen from Fig. 2

## V. Results

The methodology to extract the internal friction coefficient is illustrated using a molecule with parameters: {*b* = 800, *l*_H_ = 500 nm, *K* = 3.0 × 10^*−*9^ kg/s, *h*^∗^ = 0.0} as an example. An ensemble of such molecules is pulled from an initial trap position of 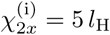 to a final trap position of 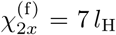 at different pulling velocities. At each value of the pulling velocity, the average dissipated work is calculated at several values of the solvent viscosity in the range, *η*_s_ = *η*_s,0_ to *η*_s_ = 10 × *η*_s,0_. In an experimental setting with water as the solvent, suitable viscogens, such as glucose or sucrose, may be added to the solvent in order to realize an approximately four-fold increase in its viscosity.^13,16^ In experiments that study the kinetics of intrachain contact formation in polypeptides^49^ suspended in a solvent mixture of ethanol and glycerol, the solvent viscosity was varied over two orders of magnitude by adjusting the proportion of glycerol in the mixture.

As shown in Fig. 3, for each value of the pulling velocity used, the average dissipated work in the hypothetical limit of zero solvent viscosity, 〈*W*_dis_〉_*η*_s_ → 0_, is obtained from a linear fit to the average dissipated work at finite solvent viscosities. In the inset of Fig. 3, the extrapolated values of the average dissipated work in the limit of zero solvent viscosity (divided by the stretching distance *d*), is plotted against the pulling velocity. The slope of the graph (*K*_BD_) represents the internal friction coefficient extracted from simulations.

**FIG. 3.**
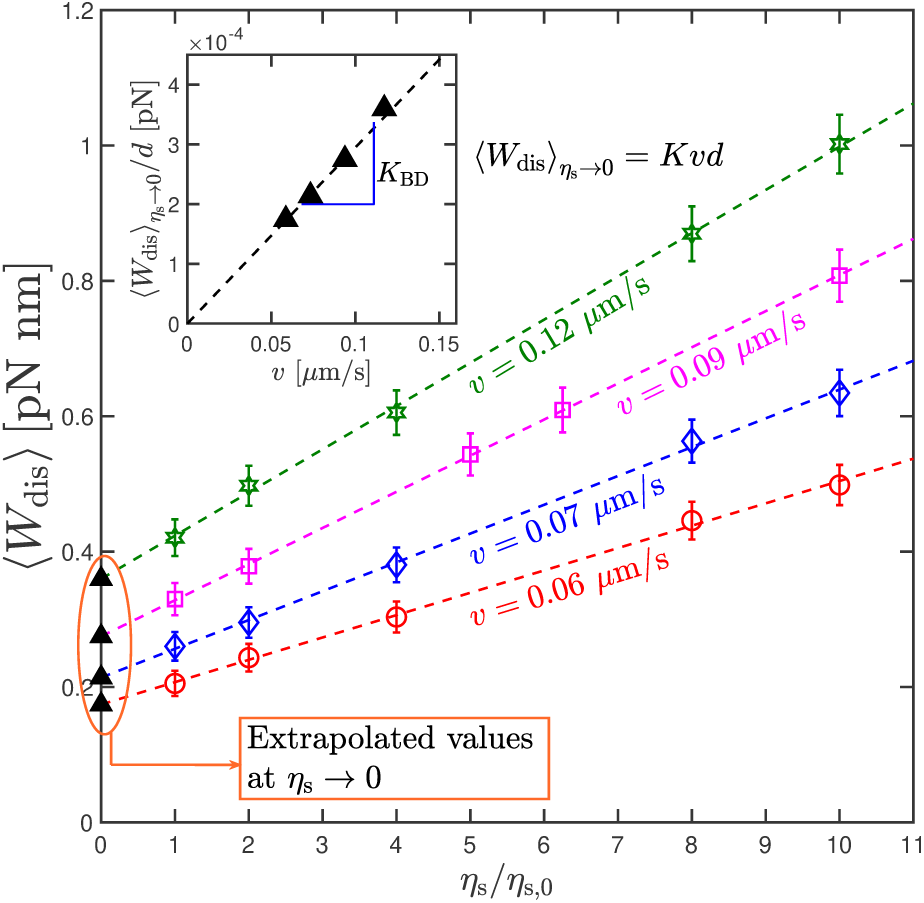
Protocol for the extraction of the internal friction coefficient. Average dissipated work as a function of the solvent viscosity, for molecules with the parameters: {*b* = 800, *l*_H_ = 500 nm, *K* = 3.0 × 10^*−*9^ kg/s}, subjected to pulling denoted by 5 *l*_H_ → 7*l*_H_ at various values of the pulling velocity, *v*, for an ensemble size, *N* = 1 × 10^4^. (Inset) The extrapolated values of 〈*W*_dis_〉 in the hypothetical limit of zero solvent viscosity, divided by the stretching distance, as a function of the pulling velocity. The slope of the graph, *K*_BD_, is an estimate of the internal friction coefficient. Error bars represent a statistical uncertainty of one standard error of the mean (s.e.m). Error bars on the extrapolated values are smaller than the symbol size.

Table I shows a comparison between the value of the internal friction coefficient used as an input parameter in the Brownian dynamics simulations, and the corresponding value extracted from the dissipated work using the protocol proposed here, for various molecular and control parameters. As seen from the table, our protocol recovers the input internal friction coefficient to within 5% accuracy. Furthermore, values of the extracted internal friction coefficient, for models with and without fluctuating hydrodynamic interactions, lie close to each other, supporting the conclusion that hydrodynamic interactions do not affect the dissipated work in the system.

**TABLE I.** Internal friction coefficients estimated using the protocol described in Fig. 3, for various values of the molecular, and control parameters. The error associated with the protocol is calculated as, % error = 100 *×* [(*K*_BD_ − *K*)*/K*].

| Parameters | Input, $K [\times 10^9 \text{ kg/s}]$ | $h^* = 0.0$ | | $h^* > 0.0$ | | |
| --- | --- | --- | --- | --- | --- | --- |
| | | $K_{BD} [\times 10^9 \text{ kg/s}]$ | % error | $h^*$ | $K_{BD} [\times 10^9 \text{ kg/s}]$ | % error |
| $b = 200, l_H = 150 \text{ nm}, 2 l_H \rightarrow 3 l_H$ | 1.0 | $1.04 \pm 0.01$ | 4.42 | 0.3 | $0.97 \pm 0.02$ | -3.11 |
| $b = 400, l_H = 350 \text{ nm}, 4 l_H \rightarrow 6 l_H$ | 1.0 | $0.99 \pm 0.03$ | -1.36 | 0.16 | $1.04 \pm 0.02$ | 4.29 |
| $b = 200, l_H = 150 \text{ nm}, 2 l_H \rightarrow 3 l_H$ | 10.0 | $10.04 \pm 0.05$ | 0.47 | 0.3 | $9.87 \pm 0.09$ | -1.24 |
| $b = 400, l_H = 350 \text{ nm}, 4 l_H \rightarrow 6 l_H$ | 10.0 | $9.922 \pm 0.008$ | -0.78 | 0.16 | $9.95 \pm 0.09$ | -0.52 |
| $b = 800, l_H = 500 \text{ nm}, 5 l_H \rightarrow 7 l_H$ | 3.0 | $2.95 \pm 0.02$ | -1.74 | — | — | — |
| $b = 800, l_H = 500 \text{ nm}, 5 l_H \rightarrow 7 l_H$ | 6.0 | $6.00 \pm 0.07$ | 0.11 | — | — | — |

## VI. Conclusion

A simple protocol for extracting the internal friction co-efficient of polymer molecules has been proposed which can be implemented experimentally by pulling the molecule multiple times using optical tweezers. The work done during the stretching process is employed to evaluate the average dissipation using the Jarzynski equality, and the dissipated work in the limit of zero solvent viscosity is then used to obtain the internal friction coefficient. Using Brownian dynamics simulations on a spring-dashpot dumbbell model for a polymer, we establish proof-of-principle by recovering the internal friction coefficient which is used as a model input. While optical tweezers offer a wide range of values from which the control parameters may be chosen, it is essential to operate in a regime with dissipation limited to a few *k_B_T*, so that: (a) the ensemble size is practically realizable, and (b) the average dissipated work is linear in the pulling velocity. We envisage that the scheme proposed here may be applicable to a variety of polymer molecules, and would enable a succinct characterisation of the dissipative properties of the molecule.

The demonstration of the proposed methodology has been made in the context of a single-mode spring-dashpot model. Though the dumbbell model offers much qualitative insight into the behavior of polymeric molecules,^31^ bead-spring-chain models with multiple springs, are needed to quantitatively describe experimental features such as the viscoelastic properties of polymer solutions^33^ and the dynamics of single molecules.^50^ Indeed the effect of internal friction on the rheology and dynamics of polymers has been studied with bead-spring-chain models, in which a dashpot with a common value of the damping coefficient is included in parallel with each of the connecting springs.^32,51,52^ It would be interesting to explore the connection between the internal friction coefficient estimated from the average dissipated work (in the limit of zero solvent viscosity) from such a model, and the damping coefficient of each dashpot in the model.

A comparison with biophysical experiments that determine the reconfiguration time of proteins, or the energy landscape of polysaccharides, would necessitate the use of a multi-bead spring chain that incorporates IV and HI. Dasbach, Manke, and Williams ^51^ have obtained an approximate analytical solution for such a chain model. The use of BD simulations to solve the bead-spring-dashpot chain model exactly is rendered difficult by the fact that formulating the correct Fokker-Planck equation for such a system, and finding the equivalent set of stochastic differential equations, is non-trivial, and is the subject of our future study.

## Acknowledgements

We thank Burkhard Dünweg, Subhashish Chaki and Ranjith Padinhateeri for enlightening discussions. The work was supported by the MonARCH and SpaceTime computational facilities of Monash University and IIT Bombay, respectively. We also acknowledge the funding and general support received from the IITB-Monash Research Academy.

