## Supplementary information for "Internal friction can be measured with the Jarzynski equality"

R. Kailasham,[1,2,3] Rajarshi Chakrabarti,[2,a)] and J. Ravi Prakash[3,b)]
[1)] *IITB-Monash Research Academy, Indian Institute of Technology Bombay, Mumbai, Maharashtra - 400076, India*
[2)] *Department of Chemistry, Indian Institute of Technology Bombay, Mumbai, Maharashtra - 400076, India*
[3)] *Department of Chemical Engineering, Monash University, Melbourne, VIC 3800, Australia*

(Dated: 16 April 2019)

The main paper presents a proof-of-concept protocol for the extraction of the internal friction coefficient of polymer molecules, which involves the use of Brownian dynamics (BD) simulations of a spring-dashpot dumbbell model. The details corresponding to various aspects of the study have been presented here.

This document is organised as follows : Sec. I presents the step-by-step formulation of the Fokker-Planck equation, and its corresponding stochastic differential equation, for the dumbbell subjected to pulling. Sec. II concerns the validation of the Jarzynski equality (JE) for our system, and compares the free energy difference obtained numerically, against those obtained from BD simulations, for two sample cases. In Sec. III, a detailed discussion of the molecular and control parameters chosen in our study is given : Sec. III A focusses on the finite extensibility parameter values and the length scales used in our study; Sec. III B discusses the effect of trap stiffness on the motion of beads, and the dissipated work, and presents a rationale for the trap stiffnesses used in our study; Sec. III C contains the force-extension profile of the spring used in our study, and analyzes the consequences of operating in the linear and non-linear regimes of this profile. Sec. III D explores the connection between the ensemble size selection for studies involving the application of the Jarzynski equality and the average dissipated work; Sec. III E compares the control parameters chosen in this study against those typically accessible in optical tweezer based pulling experiments. Additional details pertaining to the experimental feasibility of the proposed protocol is also discussed in this section. Lastly, Sec. IV gives the particulars regarding the estimation of error bars in the calculated quantities.

### I. Governing equations and solution methodology

In this work, a dumbbell model for a polymer (number of beads, $N_{\mathrm{b}}$=2) is considered, with fluctuating internal friction and hydrodynamic interactions (HI). The massless beads, each of radius $a$, are joined by a spring, with maximum stretchability $Q_0$, in parallel with a dashpot of damping coefficient $K$, and the dumbbell is suspended in an incompressible, Newtonian solvent of viscosity $\eta_{\mathrm{s}}$. The velocity field at any location $\boldsymbol{r}_{\mathrm{f}}$ in the fluid is given by $\boldsymbol{v}_{\mathrm{f}}(\boldsymbol{r},t) \equiv \boldsymbol{v}_0 + \boldsymbol{\kappa}(t)\cdot\boldsymbol{r}_{\mathrm{f}}$, where $\boldsymbol{v}_0$ is a constant vector, and $\boldsymbol{\kappa} \equiv (\nabla \boldsymbol{v}_{\mathrm{f}})^{\boldsymbol{T}}$ is the transpose of the velocity gradient tensor. In the present work, both $\boldsymbol{v}_0$ and $\boldsymbol{\kappa}$ are set to $\boldsymbol{0}$ as the pulling experiments are simulated in a quiescent fluid. However, these terms have been included in the governing equations for the sake of generality.

The two beads are manipulated by means of optical traps of stiffness $H_1 \equiv c_1 H$, and $H_2 \equiv c_2 H$, located at $\boldsymbol{\chi}_1$ and $\boldsymbol{\chi}_2$, respectively. The positions of the two beads are $\boldsymbol{r}_1$ and $\boldsymbol{r}_2$. The connector vector joining the two beads is denoted by $\boldsymbol{Q} \equiv \boldsymbol{r}_2 - \boldsymbol{r}_1$, and the centre-of-mass coordinates given by $\boldsymbol{R} \equiv (1/2)(\boldsymbol{r}_1 + \boldsymbol{r}_2)$. The configurational distribution function, $\Psi(\boldsymbol{Q},\boldsymbol{R},t)$, denotes the probability of finding the dumbbell at a position between $\boldsymbol{R}$ and $\boldsymbol{R}+d\boldsymbol{R}$, with an extension that lies between $\boldsymbol{Q}$ and $\boldsymbol{Q}+d\boldsymbol{Q}$, at any time $t$. The total restoring force in the connector vector, $\boldsymbol{F}^{(\mathrm{c})}$, has contributions from both the spring, $\boldsymbol{F}^{(\mathrm{s})} \equiv \partial U_{\mathrm{MS}}/\partial \boldsymbol{Q}$, and the dashpot, $\boldsymbol{F}^{(\mathrm{d})}$

$$\boldsymbol{F}^{(\mathrm{c})} = \boldsymbol{F}^{(\mathrm{s})} + \boldsymbol{F}^{(\mathrm{d})}, \tag{1}$$

with

$$\boldsymbol{F}^{(\mathrm{s})} = \frac{HQ_0}{3}\left[\frac{1}{2\left(1-Q/Q_0\right)^2} - \frac{1}{2} + 2\left(\frac{Q}{Q_0}\right)\right]\frac{\boldsymbol{Q}}{Q}, \tag{2}$$

where the potential energy of the spring, $U_{\mathrm{MS}}$, is given by

$$U_{\mathrm{MS}} = \frac{HQ_0^2}{3}\left[\frac{1}{2\left(1-Q/Q_0\right)} - \frac{1}{2}\left(\frac{Q}{Q_0}\right) + \left(\frac{Q}{Q_0}\right)^2\right],$$

and

$$\boldsymbol{F}^{(\mathrm{d})} = K\frac{\boldsymbol{Q}\boldsymbol{Q}}{\boldsymbol{Q}^2}\cdot [\![\dot{\boldsymbol{Q}}]\!] \tag{3}$$

Here, $K$ is the internal friction coefficient, and $[\![\dot{\boldsymbol{Q}}]\!]$ is the momentum-averaged rate-of-change of the connector vector, $\boldsymbol{Q}$.

The Fokker-Planck equation for the configurational distribution function $\Psi(\boldsymbol{Q},\boldsymbol{R},t)$ can be derived by following the procedure commonly used in polymer kinetic theory, i.e., by combining a force balance on the beads with an equation of continuity in probability space.[1] The force balance essentially states that the sum of (i) the internal friction force due to the dashpot, (ii) the restoring

[a)] Electronic mail:
[b)] Electronic mail:

force due to the finitely extensible spring, (iii) the force due to the optical traps, (iv) the random Brownian force due to bombardment by solvent molecules, and (v) the hydrodynamic force responsible for the solvent-mediated propagation of momentum on each bead, must sum up to zero.

The force balance can be solved to obtain the following equations of motion for the position vectors of the beads,[2]

$$[\![\dot{\boldsymbol{r}}_1]\!] = \left[\boldsymbol{\delta} - \frac{\epsilon\beta}{\epsilon\beta+2}\frac{\boldsymbol{QQ}}{Q^2}\right]\cdot\left(\boldsymbol{v}_0 + \boldsymbol{\kappa}\cdot\boldsymbol{r}_1 - \frac{1}{2\zeta}\boldsymbol{M}\cdot\frac{\partial U_{\mathrm{MS}}}{\partial \boldsymbol{r}_1} - \frac{k_BT}{2\zeta}\boldsymbol{M}\cdot\frac{\partial \ln\Psi}{\partial \boldsymbol{r}_1} + \frac{\epsilon}{4}\boldsymbol{M}\cdot\frac{\boldsymbol{QQ}}{Q^2}\cdot[\![\dot{\boldsymbol{r}}_2]\!] - \frac{H_1}{\zeta}(\boldsymbol{r}_1-\boldsymbol{\chi}_1) - \boldsymbol{\Omega}\cdot H_2(\boldsymbol{r}_2-\boldsymbol{\chi}_2)\right) \quad (4)$$

and

$$[\![\dot{\boldsymbol{r}}_2]\!] = \left[\boldsymbol{\delta} - \frac{\epsilon\beta}{\epsilon\beta+2}\frac{\boldsymbol{QQ}}{Q^2}\right]\cdot\left(\boldsymbol{v}_0 + \boldsymbol{\kappa}\cdot\boldsymbol{r}_2 - \frac{1}{2\zeta}\boldsymbol{M}\cdot\frac{\partial U_{\mathrm{MS}}}{\partial \boldsymbol{r}_2} - \frac{k_BT}{2\zeta}\boldsymbol{M}\cdot\frac{\partial \ln\Psi}{\partial \boldsymbol{r}_2} + \frac{\epsilon}{4}\boldsymbol{M}\cdot\frac{\boldsymbol{QQ}}{Q^2}\cdot[\![\dot{\boldsymbol{r}}_1]\!] - \frac{H_2}{\zeta}(\boldsymbol{r}_2-\boldsymbol{\chi}_2) - \boldsymbol{\Omega}\cdot H_1(\boldsymbol{r}_1-\boldsymbol{\chi}_1)\right) \quad (5)$$

where $\zeta (:= 6\pi\eta_s a)$ is the bead friction coefficient, $\epsilon := 2K/\zeta$ is the internal friction parameter, $\boldsymbol{\Omega}$ is the hydrodynamic interaction tensor, defined as

$$\boldsymbol{\Omega}(\boldsymbol{Q}) = \frac{h}{\zeta Q}\left(A\boldsymbol{\delta} + B\frac{\boldsymbol{QQ}}{Q^2}\right) \quad (6)$$

where $h = (3/4)\,a$ and $\boldsymbol{M} = 2(\boldsymbol{\delta} - \zeta\boldsymbol{\Omega})$. The terms $A$ and $B$ depend on the choice of the expression for the HI tensor, as shown in ref. 2. Here we choose the Rotne-Prager-Yamakawa (RPY) expression for the HI tensor,[3,4] in which the variables $A$ and $B$ are defined as follows

$$A = 1 + \frac{2}{3}\left(\frac{a}{Q}\right)^2 ; B = 1 - 2\left(\frac{a}{Q}\right)^2 \text{ for } Q \geq 2a \quad (7)$$

$$A = \frac{4}{3}\left(\frac{Q}{a}\right) - \frac{3}{8}\left(\frac{Q}{a}\right)^2 ; B = \frac{1}{8}\left(\frac{Q}{a}\right)^2 \text{ for } Q < 2a \quad (8)$$

The quantity $\beta$ that appears in Eqs. (4) and (5) is defined as

$$\beta = 1 - \frac{h}{Q}(A+B) \quad (9)$$

Using $\boldsymbol{r}_1 = \boldsymbol{R} - (1/2)\boldsymbol{Q}$ ; $\boldsymbol{r}_2 = \boldsymbol{R} + (1/2)\boldsymbol{Q}$, and the chain-rule for partial differentiation to operate on $\partial\Psi/\partial\boldsymbol{r}_1$ and $\partial\Psi/\partial\boldsymbol{r}_2$, leads to,

$$[\![\dot{\boldsymbol{r}}_1]\!] = \boldsymbol{v}_0 + \boldsymbol{\kappa}\cdot\left(\boldsymbol{R} - \frac{1}{2}\boldsymbol{Q}\right) + \boldsymbol{\Omega}\cdot\left(-k_BT\left[\frac{1}{2}\frac{\partial\ln\Psi}{\partial\boldsymbol{R}} + \frac{\partial\ln\Psi}{\partial\boldsymbol{Q}}\right] - \frac{\partial U_{\mathrm{MS}}}{\partial\boldsymbol{r}_2} - K\frac{\boldsymbol{QQ}}{Q^2}\cdot[\![\dot{\boldsymbol{Q}}]\!] - H_2\left(\boldsymbol{R} + \frac{1}{2}\boldsymbol{Q} - \boldsymbol{\chi}_2\right)\right) - \frac{k_BT}{\zeta}\left[\frac{1}{2}\frac{\partial\ln\Psi}{\partial\boldsymbol{R}} - \frac{\partial\ln\Psi}{\partial\boldsymbol{Q}}\right] - \frac{1}{\zeta}\frac{\partial U_{\mathrm{MS}}}{\partial\boldsymbol{r}_1} + \frac{\epsilon}{2}\frac{\boldsymbol{QQ}}{Q^2}\cdot[\![\dot{\boldsymbol{Q}}]\!] - \frac{H_1}{\zeta}\left(\boldsymbol{R} - \frac{1}{2}\boldsymbol{Q} - \boldsymbol{\chi}_1\right) \quad (10)$$

and

$$[\![\dot{\boldsymbol{r}}_2]\!] = \boldsymbol{v}_0 + \boldsymbol{\kappa}\cdot\left(\boldsymbol{R} + \frac{1}{2}\boldsymbol{Q}\right) + \boldsymbol{\Omega}\cdot\left(-k_BT\left[\frac{1}{2}\frac{\partial\ln\Psi}{\partial\boldsymbol{R}} - \frac{\partial\ln\Psi}{\partial\boldsymbol{Q}}\right] - \frac{\partial U_{\mathrm{MS}}}{\partial\boldsymbol{r}_1} + K\frac{\boldsymbol{QQ}}{Q^2}\cdot[\![\dot{\boldsymbol{Q}}]\!] - H_1\left(\boldsymbol{R} - \frac{1}{2}\boldsymbol{Q} - \boldsymbol{\chi}_1\right)\right) - \frac{k_BT}{\zeta}\left[\frac{1}{2}\frac{\partial\ln\Psi}{\partial\boldsymbol{R}} + \frac{\partial\ln\Psi}{\partial\boldsymbol{Q}}\right] - \frac{1}{\zeta}\frac{\partial U_{\mathrm{MS}}}{\partial\boldsymbol{r}_2} + \frac{\epsilon}{2}\frac{\boldsymbol{QQ}}{Q^2}\cdot[\![\dot{\boldsymbol{Q}}]\!] - \frac{H_2}{\zeta}\left(\boldsymbol{R} + \frac{1}{2}\boldsymbol{Q} - \boldsymbol{\chi}_2\right) \quad (11)$$

By adding and subtracting Eqs. (10) and (11) suitably, we obtain

$$[\![\dot{\boldsymbol{R}}]\!] = \boldsymbol{v}_0 + \boldsymbol{\kappa}\cdot\boldsymbol{R} - \frac{k_BT}{2\zeta}(\boldsymbol{\delta} + \zeta\boldsymbol{\Omega})\cdot\frac{\partial\ln\Psi}{\partial\boldsymbol{R}} - \frac{1}{2\zeta}(\boldsymbol{\delta} + \zeta\boldsymbol{\Omega})\cdot\boldsymbol{X} \quad (12)$$

$$[\![\dot{\boldsymbol{Q}}]\!] = \left[\boldsymbol{\delta} - \frac{\epsilon\beta}{\epsilon\beta+1}\frac{\boldsymbol{QQ}}{Q^2}\right]\cdot\left(\boldsymbol{\kappa}\cdot\boldsymbol{Q} - \frac{k_BT}{\zeta}\boldsymbol{M}\cdot\frac{\partial}{\partial\boldsymbol{Q}}\ln\Psi - \frac{1}{\zeta}\boldsymbol{M}\cdot\frac{\partial U_{\mathrm{MS}}}{\partial\boldsymbol{Q}} - \frac{1}{2\zeta}\boldsymbol{M}\cdot\boldsymbol{Y}\right) \quad (13)$$

where

$$\boldsymbol{X} = \boldsymbol{R}\left(H_2 + H_1\right) + \boldsymbol{Q}\left(\frac{H_2 - H_1}{2}\right) - \left(H_2\boldsymbol{\chi}_2 + H_1\boldsymbol{\chi}_1\right)$$

and

$$\boldsymbol{Y} = \boldsymbol{R}\left(H_2 - H_1\right) + \boldsymbol{Q}\left(\frac{H_2 + H_1}{2}\right) - \left(H_2\boldsymbol{\chi}_2 - H_1\boldsymbol{\chi}_1\right)$$

and both $\boldsymbol{X}$ and $\boldsymbol{Y}$ have dimensions of force.

The equation of continuity in terms of $\boldsymbol{R}$ and $\boldsymbol{Q}$ is given by,[1]

$$\frac{\partial\Psi}{\partial t} = -\left(\frac{\partial}{\partial\boldsymbol{R}}\cdot[\![\dot{\boldsymbol{R}}]\!]\Psi\right) - \left(\frac{\partial}{\partial\boldsymbol{Q}}\cdot[\![\dot{\boldsymbol{Q}}]\!]\Psi\right) \tag{14}$$

Substituting Eqs. (12) and (13) into the above expression leads to the Fokker-Planck equation that governs the configurational distribution function $\Psi\left(\boldsymbol{Q},\boldsymbol{R},t\right)$,

$$\begin{aligned}\frac{\partial\Psi}{\partial t} &= -\frac{\partial}{\partial\boldsymbol{R}}\cdot\left\{\left[\boldsymbol{v}_0 + \boldsymbol{\kappa}\cdot\boldsymbol{R} - \frac{1}{2\zeta}\left(\boldsymbol{\delta}+\zeta\boldsymbol{\Omega}\right)\cdot\boldsymbol{X}\right]\Psi\right\} + \frac{k_BT}{2\zeta}\frac{\partial}{\partial\boldsymbol{R}}\cdot\left(\boldsymbol{\delta}+\zeta\boldsymbol{\Omega}\right)\cdot\frac{\partial\Psi}{\partial\boldsymbol{R}} \\ &\quad - \frac{\partial}{\partial\boldsymbol{Q}}\cdot\left\{\left[\left[\boldsymbol{\delta} - \frac{\epsilon\beta}{\epsilon\beta+1}\frac{\boldsymbol{Q}\boldsymbol{Q}}{Q^2}\right]\cdot\left(\boldsymbol{\kappa}\cdot\boldsymbol{Q} - \frac{1}{\zeta}\boldsymbol{M}\cdot\frac{\partial U_{\rm MS}}{\partial\boldsymbol{Q}} - \frac{1}{2\zeta}\boldsymbol{M}\cdot\boldsymbol{Y}\right)\right]\Psi\right\} \\ &\quad + \frac{k_BT}{\zeta}\frac{\partial}{\partial\boldsymbol{Q}}\cdot\left[\left(\boldsymbol{\delta} - \frac{\epsilon\beta}{\epsilon\beta+1}\frac{\boldsymbol{Q}\boldsymbol{Q}}{Q^2}\right)\cdot\boldsymbol{M}\right]\cdot\frac{\partial\Psi}{\partial\boldsymbol{Q}}\end{aligned} \tag{15}$$

Introducing the length scale, $l_{\rm H} = \sqrt{k_BT/H}$, the time scale $\lambda_{\rm H} = \zeta/4H$, and scaling units of energy by $k_BT$, and those of force by $\sqrt{k_BTH}$, the following dimensionless quantities (denoted with an asterisk as a superscript), can be defined,

$$t^* = \frac{t}{\lambda_{\rm H}};\ \boldsymbol{Q}^* = \frac{\boldsymbol{Q}}{l_{\rm H}};\ b = \frac{Q_0^2}{l_{\rm H}^2};\ \boldsymbol{\kappa}^* = \lambda_{\rm H}\boldsymbol{\kappa};\ U^*_{\rm MS} = \frac{U_{\rm MS}}{k_BT};\ \Psi^* = \Psi l_{\rm H}^3;\ \boldsymbol{X}^* = \frac{\boldsymbol{X}}{\sqrt{k_BTH}} \tag{16}$$

In terms of these non-dimensional variables, the Fokker-Planck equation assumes the following form,

$$\begin{aligned}\frac{\partial\Psi^*}{\partial t^*} &= -\frac{\partial}{\partial\boldsymbol{R}^*}\cdot\left\{\left[\boldsymbol{v}_0^* + \boldsymbol{\kappa}^*\cdot\boldsymbol{R}^* - \frac{1}{8}\left(\boldsymbol{\delta}+\zeta\hat{\boldsymbol{\Omega}}\right)\cdot\boldsymbol{X}^*\right]\Psi^*\right\} + \frac{1}{8}\frac{\partial}{\partial\boldsymbol{R}}\cdot\left(\boldsymbol{\delta}+\zeta\hat{\boldsymbol{\Omega}}\right)\cdot\frac{\partial\Psi^*}{\partial\boldsymbol{R}^*} \\ &\quad - \frac{\partial}{\partial\boldsymbol{Q}^*}\cdot\left\{\left[\left[\boldsymbol{\delta} - \frac{\epsilon\beta^*}{\epsilon\beta^*+1}\frac{\boldsymbol{Q}^*\boldsymbol{Q}^*}{Q^{*2}}\right]\cdot\left(\boldsymbol{\kappa}^*\cdot\boldsymbol{Q}^* - \frac{1}{4}\boldsymbol{M}^*\cdot\frac{\partial U^*_{\rm MS}}{\partial\boldsymbol{Q}^*} - \frac{1}{8}\boldsymbol{M}^*\cdot\boldsymbol{Y}^*\right)\right]\Psi^*\right\} \\ &\quad + \frac{1}{4}\frac{\partial}{\partial\boldsymbol{Q}^*}\cdot\left[\left(\boldsymbol{\delta} - \frac{\epsilon\beta^*}{\epsilon\beta^*+1}\frac{\boldsymbol{Q}^*\boldsymbol{Q}^*}{Q^{*2}}\right)\cdot\boldsymbol{M}^*\right]\cdot\frac{\partial\Psi^*}{\partial\boldsymbol{Q}^*}\end{aligned} \tag{17}$$

where

$$\hat{\boldsymbol{\Omega}}\left(\boldsymbol{Q}^*\right) = \frac{\alpha}{\zeta Q^*}\left(A^*\boldsymbol{\delta} + B^*\frac{\boldsymbol{Q}^*\boldsymbol{Q}^*}{Q^{*2}}\right) \tag{18}$$

with $\alpha = (3/4)\sqrt{\pi}h^*$ and $h^* = a/(\sqrt{\pi}l_{\rm H})$. The quantity $\beta^*$ is dimensionless and is defined as,

$$\beta^* = 1 - \frac{\alpha}{Q^*}\left(A^* + B^*\right) \tag{19}$$

Using the following identity,

$$\frac{\partial}{\partial\boldsymbol{x}}\cdot\left[\boldsymbol{L}\cdot\frac{\partial f}{\partial\boldsymbol{x}}\right] = \frac{\partial}{\partial\boldsymbol{x}}\frac{\partial}{\partial\boldsymbol{x}}:\left[\boldsymbol{L}^{\rm T}f\right] - \frac{\partial}{\partial\boldsymbol{x}}\cdot\left[f\frac{\partial}{\partial\boldsymbol{x}}\cdot\boldsymbol{L}^{\rm T}\right] \tag{20}$$

where $\boldsymbol{L}$ is a tensor and $f$ is a scalar, the second and fourth terms on the right-hand-side of Eq. (17) can rewritten in a way that renders the Fokker-Planck equation amenable to Itô's interpretation. Since $\hat{\boldsymbol{\Omega}}\left(\boldsymbol{Q}^*\right)$ is independent of $\boldsymbol{R}^*$, and

$$\left(\boldsymbol{\delta} - \frac{\epsilon\beta^*}{\epsilon\beta^*+1}\frac{\boldsymbol{Q}^*\boldsymbol{Q}^*}{Q^{*2}}\right)\cdot\left(\boldsymbol{\delta} - \zeta\hat{\boldsymbol{\Omega}}\right) = \left(\frac{Q^* - A^*\alpha}{Q^*}\right)\times\left(\boldsymbol{\delta} - g_1\frac{\boldsymbol{Q}^*\boldsymbol{Q}^*}{Q^{*2}}\right)$$

the Fokker-Planck equation can be rewritten as follows,

$$\frac{\partial \Psi^*}{\partial t^*} = -\frac{\partial}{\partial \boldsymbol{R^*}} \cdot \left\{ \left[ \boldsymbol{v}_0^* + \boldsymbol{\kappa}^* \cdot \boldsymbol{R}^* - \frac{1}{8}\left(\boldsymbol{\delta} + \zeta \hat{\boldsymbol{\Omega}}\right) \cdot \boldsymbol{X}^* \right] \Psi^* \right\} + \frac{1}{2}\frac{\partial}{\partial \boldsymbol{R^*}}\frac{\partial}{\partial \boldsymbol{R^*}} : \left[ \frac{\left(\boldsymbol{\delta} + \zeta \hat{\boldsymbol{\Omega}}\right)}{4} \Psi^* \right]$$
$$- \frac{\partial}{\partial \boldsymbol{Q^*}} \cdot \left\{ \left[ \frac{g_2}{2}\frac{\boldsymbol{Q^*}}{Q^*} + \left[ \boldsymbol{\delta} - \frac{\epsilon\beta^*}{\epsilon\beta^* + 1}\frac{\boldsymbol{Q^*Q^*}}{Q^{*2}} \right] \cdot \left( \boldsymbol{\kappa}^* \cdot \boldsymbol{Q}^* - \frac{1}{4}\boldsymbol{M}^* \cdot \frac{\partial U^*_{\text{MS}}}{\partial \boldsymbol{Q^*}} - \frac{1}{8}\boldsymbol{M}^* \cdot \boldsymbol{Y}^* \right) \right] \Psi^* \right\}$$
$$+ \frac{1}{2}\frac{\partial}{\partial \boldsymbol{Q^*}}\frac{\partial}{\partial \boldsymbol{Q^*}} : \left[ \left( \frac{Q^* - A^*\alpha}{Q^*} \right) \left( \boldsymbol{\delta} - g_1 \frac{\boldsymbol{Q^*Q^*}}{Q^{*2}} \right) \Psi^* \right] \tag{21}$$

where

$$g_1 = \frac{\alpha B^* Q^* + \epsilon(Q^* - A^*\alpha)[Q^* - \alpha(A^* + B^*)]}{(Q^* - A^*\alpha)\left\{Q^* + \epsilon[Q^* - \alpha(A^* + B^*)]\right\}} \tag{22}$$

$$g_2 = \frac{2\alpha B^*}{\left\{Q^* + \epsilon[Q^* - \alpha(A^* + B^*)]\right\}^2} - 2g_1\left(\frac{Q^* - A^*\alpha}{Q^{*2}}\right)$$

The Fokker-Planck equation (Eq. (21)) can be written in the following compact form,

$$\frac{\partial \Psi^*}{\partial t^*} = -\frac{\partial}{\partial \boldsymbol{R^*}} \cdot \left\{ \boldsymbol{e}\Psi^* \right\} + \frac{1}{2}\frac{\partial}{\partial \boldsymbol{R^*}}\frac{\partial}{\partial \boldsymbol{R^*}} : [\boldsymbol{E}\Psi^*]$$
$$- \frac{\partial}{\partial \boldsymbol{Q^*}} \cdot \left\{ \boldsymbol{g}\Psi^* \right\} + \frac{1}{2}\frac{\partial}{\partial \boldsymbol{Q^*}}\frac{\partial}{\partial \boldsymbol{Q^*}} : [\boldsymbol{G}\Psi^*] \tag{23}$$

where the definitions of the quantities $\boldsymbol{e}, \boldsymbol{E}, \boldsymbol{g}$ and $\boldsymbol{G}$ are clear by comparison of Eqs. (21) and (23). It is convenient to define a collective variable, $\boldsymbol{C}$, which is a six-element vector containing the components of $\boldsymbol{R^*}$ and $\boldsymbol{Q^*}$, such that $\boldsymbol{C} \equiv \left[R^*_x, R^*_y, R^*_z, Q^*_x, Q^*_y, Q^*_z\right]$. Similarly, a six-element vector $\boldsymbol{j}$ can be defined, containing the components of $\boldsymbol{e}$ and $\boldsymbol{g}$, along with the definition of a $2 \times 2$ block matrix $\boldsymbol{D}$, whose off-diagonal elements are $\boldsymbol{0}$, and the diagonal elements are the matrices $\boldsymbol{E}$ and $\boldsymbol{G}$ (each of size $3 \times 3$). With these definitions, the Fokker-Planck equation in Eq. (23) can be written as

$$\frac{\partial \Psi^*}{\partial t^*} = -\frac{\partial}{\partial \boldsymbol{C}} \cdot \left\{ \boldsymbol{j}\Psi^* \right\} + \frac{1}{2}\frac{\partial}{\partial \boldsymbol{C}}\frac{\partial}{\partial \boldsymbol{C}} : [\boldsymbol{D}\Psi^*] \tag{24}$$

The stochastic differential equation (SDE) corresponding to Eq. (24) can be obtained using the Itô interpretation, as

$$d\boldsymbol{C} = \boldsymbol{j}dt^* + \boldsymbol{b} \cdot d\boldsymbol{w}_t \tag{25}$$

where $\boldsymbol{w}_t$ is a Wiener process and $\boldsymbol{b} \cdot \boldsymbol{b}^T = \boldsymbol{D}$. The SDE (Eq. (25)) is solved using a semi-implicit predictor-corrector scheme,[5] as discussed in the following subsection.

#### Solver details

With reference to Eq. (24), $\boldsymbol{D}$ is a $6 \times 6$ matrix, and its square root, $\boldsymbol{b}_j$, at any time $t^*_j$ is found using Cholesky decomposition.[6] Although Eq. (25) is written in terms of the collective variable $\boldsymbol{C}$, the equation for $\boldsymbol{R^*}$ is solved purely explicitly, whereas the equation in $\boldsymbol{Q^*}$ is solved by treating only the spring force term implicitly. For the sake of clarity, the predictor and corrector equations for $\boldsymbol{R^*}$ and $\boldsymbol{Q^*}$ are presented separately. It is useful to define another six-element vector, $\Delta\boldsymbol{S}_j$, as

$$\Delta\boldsymbol{S}_j = \boldsymbol{b}_j \cdot \Delta\boldsymbol{w}_j \tag{26}$$

where $\boldsymbol{w}_j$ is a vector of six independent Weiner processes, each of mean zero and variance $\Delta t^*_j$. The first three elements of $\Delta\boldsymbol{S}_j$, denoted by $\Delta\boldsymbol{S}_j^{(R)}$, contain the noise contribution to $\boldsymbol{R}^*$, and the next three elements, denoted by $\Delta\boldsymbol{S}_j^{(Q)}$, contribute to the noise in $\boldsymbol{Q^*}$. In the following discussion, Eqs. (27)—(34) are in their dimensionless form, but the asterisk has been dropped from these equations for the sake of notational simplicity.

### Predictor step

$$\tilde{\boldsymbol{R}}(t_{j+1}) = \boldsymbol{R}(t_j) + \left[\boldsymbol{v}_0 + \boldsymbol{\kappa}(t_j) \cdot \boldsymbol{R}(t_j) - \boldsymbol{X}_a(t_j)\right]\Delta t_j + \Delta\boldsymbol{S}_j^{(R)} \tag{27}$$

$$\tilde{\boldsymbol{Q}}\left(t_{j+1}\right)=\boldsymbol{Q}(t_j)+\Bigg[\boldsymbol{\kappa}(t_j)\cdot\boldsymbol{Q}(t_j)-\left(\frac{\epsilon\beta(t_j)}{\epsilon\beta(t_j)+1}\right)\left[\boldsymbol{\kappa}(t_j):\frac{\boldsymbol{Q}(t_j)\boldsymbol{Q}(t_j)}{Q^2(t_j)}\right]\boldsymbol{Q}(t_j)$$
$$-\frac{f}{2}\left(\frac{\beta(t_j)}{\epsilon\beta(t_j)+1}\right)\frac{\boldsymbol{Q}(t_j)}{Q(t_j)}+\frac{g_2(t_j)}{2}\frac{\boldsymbol{Q}(t_j)}{Q(t_j)}-\boldsymbol{Y}_a(t_j)\Bigg]\Delta t_j+\Delta\boldsymbol{S}_j^{(Q)} \tag{28}$$

where

$$\boldsymbol{X}_a(t_j)=\frac{1}{8}\left(\boldsymbol{\delta}+\zeta\hat{\boldsymbol{\Omega}}(\boldsymbol{Q}_j)\right)\cdot\boldsymbol{X}\left(\boldsymbol{Q}_j,\boldsymbol{R}_j\right)$$
$$\boldsymbol{Y}_a(t_j)=\frac{1}{4}\left(\boldsymbol{\delta}-\frac{\epsilon\beta(t_j)}{\epsilon\beta(t_j)+1}\frac{\boldsymbol{Q}(t_j)\boldsymbol{Q}(t_j)}{Q^2(t_j)}\right)\cdot\left(\boldsymbol{\delta}-\zeta\hat{\boldsymbol{\Omega}}(\boldsymbol{Q}_j)\right)\cdot\boldsymbol{Y}\left(\boldsymbol{Q}_j,\boldsymbol{R}_j\right) \tag{29}$$
$$f(t_j)=\frac{\sqrt{b}}{3}\left[\frac{1}{2\left(1-Q(t_j)/\sqrt{b}\right)^2}-\frac{1}{2}+2\left(\frac{Q(t_j)}{\sqrt{b}}\right)\right]$$

and the notations $\boldsymbol{Q}_j$ and $\boldsymbol{Q}(t_j)$ have been used interchangeably to refer to the same quantity.

**Corrector step**

$$\boldsymbol{R}(t_{j+1})=\tilde{\boldsymbol{R}}\left(t_{j+1}\right)+\frac{1}{2}\left[\boldsymbol{\kappa}(t_{j+1})\cdot\tilde{\boldsymbol{R}}_{j+1}-\boldsymbol{\kappa}(t_j)\cdot\tilde{\boldsymbol{R}}_j-\tilde{\boldsymbol{X}}_a(t_{j+1})+\boldsymbol{X}_a(t_j)\right]\Delta t_j \tag{30}$$

$$\left[1+\frac{f(t_{j+1})}{4Q(t_{j+1})}\left(\frac{\tilde{\beta}(t_{j+1})}{\epsilon\tilde{\beta}(t_{j+1})+1}\right)\Delta t_j\right]\boldsymbol{Q}(t_{j+1})=\tilde{\boldsymbol{Q}}\left(t_{j+1}\right)$$
$$+\frac{1}{2}\left[\boldsymbol{\kappa}(t_{j+1})\cdot\tilde{\boldsymbol{Q}}_{j+1}-\boldsymbol{\kappa}(t_j)\cdot\tilde{\boldsymbol{Q}}_j+\tilde{\boldsymbol{q}}(t_{j+1})-\boldsymbol{q}(t_j)-\tilde{\boldsymbol{Y}}_a(t_{j+1})+\boldsymbol{Y}_a(t_j)\right]\Delta t_j \tag{31}$$

where

$$\tilde{\boldsymbol{X}}_a(t_{j+1})=\frac{1}{8}\left(\boldsymbol{\delta}+\zeta\hat{\boldsymbol{\Omega}}(\tilde{\boldsymbol{Q}}_{j+1})\right)\cdot\tilde{\boldsymbol{X}}\left(\tilde{\boldsymbol{Q}}_{j+1},\tilde{\boldsymbol{R}}_{j+1}\right)$$
$$\tilde{\boldsymbol{Y}}_a(t_{j+1})=\frac{1}{4}\left(\boldsymbol{\delta}-\frac{\epsilon\tilde{\beta}(t_{j+1})}{\epsilon\tilde{\beta}(t_{j+1})+1}\frac{\tilde{\boldsymbol{Q}}(t_{j+1})\tilde{\boldsymbol{Q}}(t_{j+1})}{\tilde{Q}^2(t_{j+1})}\right)\cdot\left(\boldsymbol{\delta}-\zeta\hat{\boldsymbol{\Omega}}(\tilde{\boldsymbol{Q}}_{j+1})\right)\cdot\boldsymbol{Y}\left(\tilde{\boldsymbol{Q}}_{j+1},\tilde{\boldsymbol{R}}_{j+1}\right)$$
$$f(t_{j+1})=\frac{\sqrt{b}}{3}\left[\frac{1}{2\left(1-Q(t_{j+1})/\sqrt{b}\right)^2}-\frac{1}{2}+2\left(\frac{Q(t_{j+1})}{\sqrt{b}}\right)\right] \tag{32}$$
$$\boldsymbol{q}(t_j)=\frac{g_2(t_j)}{2}\frac{\boldsymbol{Q}(t_j)}{Q(t_j)}-\left(\frac{\epsilon\beta(t_j)}{\epsilon\beta(t_j)+1}\right)\left[\boldsymbol{\kappa}(t_j):\frac{\boldsymbol{Q}(t_j)\boldsymbol{Q}(t_j)}{Q^2(t_j)}\right]\boldsymbol{Q}(t_j)$$
$$\tilde{\boldsymbol{q}}(t_{j+1})=\frac{g_2(t_{j+1})}{2}\frac{\tilde{\boldsymbol{Q}}(t_{j+1})}{\tilde{Q}(t_{j+1})}-\left(\frac{\epsilon\tilde{\beta}(t_{j+1})}{\epsilon\tilde{\beta}(t_{j+1})+1}\right)\left[\boldsymbol{\kappa}(t_{j+1}):\frac{\tilde{\boldsymbol{Q}}(t_{j+1})\tilde{\boldsymbol{Q}}(t_{j+1})}{\tilde{Q}^2(t_{j+1})}\right]\tilde{\boldsymbol{Q}}(t_{j+1})$$

By setting the length of the vector on the RHS of Eq. (31) to be $L$, and the length of $\boldsymbol{Q}(t_{j+1})$ to be $\Phi$, the following cubic equation is obtained:

$$\phi^3-\phi^2\left[\frac{3(3\omega+4+2\sigma)}{2(2\omega+3)}\right]+\phi\left[\frac{3(1+\omega+2\sigma)}{2\omega+3}\right]-\frac{3\sigma}{2\omega+3}=0 \tag{33}$$

where

$$\omega=\left(\frac{\tilde{\beta}(t_{j+1})}{\epsilon\tilde{\beta}(t_{j+1})+1}\right)\frac{\Delta t_j}{4}\,;\quad \phi=\frac{\Phi}{\sqrt{b}}\,;\quad \sigma=\frac{L}{\sqrt{b}} \tag{34}$$

Eq. (33) has three roots—two complex and one real—and the real root is obtained using the Newton-Raphson scheme.[6] Note that the equations are solved in their dimensionless form, and the dimensional quantities are obtained by a suitable multiplication with the scaling factors, as explained in the discussion surrounding Eq. (16).

### A. Simulation details

To begin with, the initial values of $\boldsymbol{Q}^*$ and $\boldsymbol{R}^*$ are picked from a Gaussian distribution of zero mean and unit variance. With the first trap held at the origin, and the second at $\boldsymbol{\chi}_2^{(\mathrm{i})*} = (\chi_{2x}^{(\mathrm{i})*}, 0, 0)$, the dumbbell is equilibrated for a duration of fifty-five dimensionless times. Equilibration is ascertained by checking that $\langle Q^{*2} \rangle$ has reached a steady value with respect to time. Then, the pulling is commenced (at $t^* = 0$), by varying the position of the second trap linearly, as $\chi_{2x}^* = \chi_{2x}^{(\mathrm{i})*} + v_x^* t^*$, till $t^* = \tau^*$. The window $[0, \tau^*]$ is uniformly divided into $N_\mathrm{t}$ intervals, such that $\Delta t_j^* \equiv t_j^* - t_{j-1}^* = \tau^*/N_\mathrm{t}$, where $j = 1, 2, ..., (N_\mathrm{t} + 1)$. For representative values of the molecular and control parameters, the average dissipated work is computed using the time-step widths $\Delta t^* = \{10^{-3}, 10^{-4}, 10^{-5}\}$. The results for all the time-steps concur within statistical error bars of the simulation, and the largest of the three time-step widths, i.e; $\Delta t^* = 1 \times 10^{-3}$, is used for all the cases where $c_1 = 1000$. For $c_1 = 100$, $\Delta t^* = 1 \times 10^{-2}$ is found to suffice, whereas $c_1 = 10000$ requires $\Delta t^* = 1 \times 10^{-4}$.

The dimensionless equivalent of the work done during one realization of the pulling event is calculated as follows,

$$W^* = c_2 \sum_{j=1}^{N_\mathrm{t}} \left(\chi_{2x}^* - r_{2x}^*\right)_j v_x^* \Delta t_j^* \tag{35}$$

where the subscript $j$ on the first term indicates that it is evaluated at time, $t_j^*$, and $r_{2x}^*$ refers to the $x$-coordinate of the position of the dumbbell bead subjected to pulling.

The protocol proposed here involves pulling the molecule over a pre-determined distance at the same dimensional velocity but different solvent viscosities. In this context, it is essential to note that the timescale varies linearly with the solvent viscosity, $\lambda_\mathrm{H} \propto \eta_\mathrm{s}$. In order to maintain the same dimensional pulling time ($\tau = \tau^* \lambda_\mathrm{H}$) across simulations with differing solvent viscosity, the dimensionless pulling time ($\tau^*$) is scaled by $1/\eta_\mathrm{s}$ as the solvent viscosity is increased.

### II. Validation of Jarzynski's equality

The total Hamiltonian of the dumbbell and trap system is written as

$$\mathcal{H}^* \equiv \frac{\mathcal{H}}{k_B T} = U_\mathrm{MS}^* + \frac{c_1}{2}\left(\boldsymbol{r}_1^* - \boldsymbol{\chi}_1^*\right)^2 + \frac{c_2}{2}\left(\boldsymbol{r}_2^* - \boldsymbol{\chi}_2^*\right)^2 \tag{36}$$

The expression for $\mathcal{H}^*$ can be rewritten in terms of $\boldsymbol{Q}^*$ and $\boldsymbol{R}^*$ as,

$$\begin{aligned}\mathcal{H}^* = \frac{b}{3}\left[\frac{1}{2\left(1 - Q^*/\sqrt{b}\right)} - \frac{1}{2}\left(\frac{Q^*}{\sqrt{b}}\right) + \left(\frac{Q^*}{\sqrt{b}}\right)^2\right] \\ - \boldsymbol{R}^* \cdot \left(c_1 \boldsymbol{\chi}_1^* + c_2 \boldsymbol{\chi}_2^*\right) + \frac{c_1 \chi_1^{*2} + c_2 \chi_2^{*2}}{2} \\ + \frac{\boldsymbol{Q}^*}{2} \cdot \left(c_1 \boldsymbol{\chi}_1^* - c_2 \boldsymbol{\chi}_2^*\right) - \left(\frac{c_1 - c_2}{2}\right) \boldsymbol{Q}^* \cdot \boldsymbol{R}^* \\ + R^{*2}\left(\frac{c_1 + c_2}{2}\right) + \frac{Q^{*2}}{4}\left(\frac{c_1 + c_2}{2}\right)\end{aligned} \tag{37}$$

The steady-state configurational distribution function can be written as

$$\Psi^*(\boldsymbol{Q}^*, \boldsymbol{R}^*) = \frac{1}{\mathcal{Z}} \exp[-\mathcal{H}^*] \tag{38}$$

where $\mathcal{Z}$ is the partition function of the system, given by

$$\mathcal{Z} = \int\int \exp[-\mathcal{H}^*] d\boldsymbol{R}^* d\boldsymbol{Q}^* \tag{39}$$

Substituiting the definition of $\mathcal{H}^*$ from Eq. (37) into Eq. (39) yields the following equation,

$$\mathcal{Z} = \int\left[\int \exp\left[-\bar{c}\left(\boldsymbol{R}^* \cdot \boldsymbol{R}^*\right) - m\left(\boldsymbol{R}^* \cdot \boldsymbol{l}\right)\right] d\boldsymbol{R}^*\right] \exp\left[\Xi\right] d\boldsymbol{Q}^* \tag{40}$$

where

$$\bar{c} = \frac{c_1 + c_2}{2}\,; \quad m = -1\,; \boldsymbol{l} = -\left[c_1 \boldsymbol{\chi}_1^* + c_2 \boldsymbol{\chi}_2^* + \left(\frac{c_1 - c_2}{2}\right)\right];$$

$$\Xi = -\frac{Q^{*2}}{4}\left(\frac{c_1 + c_2}{2}\right) - \frac{\boldsymbol{Q}^*}{2} \cdot \left(c_1 \boldsymbol{\chi}_1^* - c_2 \boldsymbol{\chi}_2^*\right) - \frac{c_1 \chi_1^{*2} + c_2 \chi_2^{*2}}{2} - U_\mathrm{MS}^*$$

The inner integral in Eq. (40) can be evaluated using the following identity[1] for Gaussian integrals,

$$\int \exp\left[-\bar{c}\,(\boldsymbol{u}\cdot\boldsymbol{u}) - m\,(\boldsymbol{u}\cdot\boldsymbol{j})\right] d\boldsymbol{u} = \left(\frac{\pi}{\bar{c}}\right)^{3/2} \exp\left[\frac{\nu^2}{4\bar{c}}\,(\boldsymbol{j}\cdot\boldsymbol{j})\right] \tag{41}$$

resulting in

$$\mathcal{Z} = \left(\frac{2\pi}{c_1+c_2}\right)^{3/2} \int \exp\left[\frac{\boldsymbol{l}\cdot\boldsymbol{l}}{4\bar{c}} + \Xi\right] d\boldsymbol{Q^*} \tag{42}$$

Upon simplification, one obtains

$$\mathcal{Z} = \left(\frac{2\pi}{c_1+c_2}\right)^{3/2} \int \exp\left\{-k\left[\boldsymbol{Q^*} - \boldsymbol{s^*}\right]^2 - U^*_{\text{MS}}\right\} d\boldsymbol{Q^*} \tag{43}$$

where $k = (c_1 c_2)/2(c_1+c_2)$, and $\boldsymbol{s^*} = \boldsymbol{\chi}_2^* - \boldsymbol{\chi}_1^*$. The integral in Eq. (43) can be evaluated by converting to spherical co-ordinates, recognising that $Q_x^* = Q^* \sin\theta\cos\phi$, $Q_y^* = Q^*\sin\theta\sin\phi$, $Q_z^* = Q^*\cos\theta$. Therefore,

$$\begin{aligned}\mathcal{Z} = \left(\frac{2\pi}{c_1+c_2}\right)^{3/2} \int_{Q^*=0}^{\sqrt{b}} \int_{\theta=0}^{\pi} \int_{\phi=0}^{2\pi} \Bigg[ &\exp\Big(-k\,[Q_x^* - s_x^*]^2\Big) \exp\Big(-k\,[Q_y^* - s_y^*]^2\Big) \exp\Big(-k\,[Q_z^* - s_z^*]^2\Big) \\ &\times \exp\left\{\frac{b}{3}\left[\frac{1}{2}\left(\frac{Q^*}{\sqrt{b}}\right) - \frac{1}{2\left(1 - Q^*/\sqrt{b}\right)} - \left(\frac{Q^*}{\sqrt{b}}\right)^2\right]\right\}\Bigg] Q^{*2} dQ^* \sin\theta\, d\theta\, d\phi\end{aligned} \tag{44}$$

The integral in Eq. (44) does not have an analytically closed-form solution, and is evaluated numerically using MATLAB. Using the relation between the free-energy and the partition function, $A^* = -\ln\mathcal{Z}$, it follows that the free-energy difference in going from the initial state to the final state is given by,

$$\Delta A^*_{\text{num}} = \ln\left[\frac{\mathcal{Z}(\boldsymbol{\chi}_2^* = \boldsymbol{\chi}_2^{(\text{i})*})}{\mathcal{Z}(\boldsymbol{\chi}_2^* = \boldsymbol{\chi}_2^{(\text{f})*})}\right] \tag{45}$$

where the subscript 'num' indicates that the free energy difference has been calculated numerically.

Fig. 1 shows a comparison between the free energy difference obtained from Brownian dynamics simulations of $N = 1\times 10^5$ trajectories using Jarzynski's equality,[7] and that obtained from Eq. (45), for the two parameter sets indicated in Table I.

TABLE I. Parameter values for the two representative cases for which the free energy differences are evaluated using Jarzynski's equality and numerical integration.

| | Parameter sets | |
|---|---|---|
| | 1 | 2 |
| $b$ | 50 | 80 |
| $c_1$ | 20 | 15 |
| $c_2$ | 1 | 15 |
| $\boldsymbol{\chi}_1^*$ | $(0,0,0)$ | $(0,0,0)$ |
| $\boldsymbol{\chi}_2^{(\text{i})*}$ | $(1,0,0)$ | $(4,0,0)$ |
| $\boldsymbol{\chi}_2^{(\text{f})*}$ | $(3,0,0)$ | $(5,0,0)$ |

There is excellent agreement between the results obtained using the two approaches. Furthermore, it is seen that the internal friction parameter does not affect the free energy difference at low pulling velocities. At high values of the pulling velocity, however, the free energy differences calculated for the cases with and without internal friction differ slightly. This discrepancy at higher velocities, however, is an inherent feature of Jarzynski's recipe, wherein the distribution of work values become broader at higher values of the dissipated work. The use of a higher number of trajectories for the high velocity cases could result in a better agreement between the free energy differences for cases with and without internal viscosity. The concurrence in results is also represented in tabular form (see Table II). Note that only the data corresponding to the case $\epsilon = 0.0$, from Fig. 1, has been used for the quantification of errors in Table II.

TABLE II. A comparison of the free-energy differences calculated using numerical integration [Eq. (45)], and BD simulations using Jarzynski's equality [Eq. (55)], over $N = 1 \times 10^5$ trajectories. Simulation data reported for freely-draining dumbbells with no internal friction [$h^* = 0.0$, $\epsilon = 0.0$]. The error is quantified as, $\%\,\text{error} = 100 \times [(\Delta A^* - \Delta A^*_{\text{num}})/\Delta A^*_{\text{num}}]$.

| | Parameter set 1 : | $\Delta A^*_{\text{num}} = 2.11504$ | |
|---|---|---|---|
| $v^*$ | $\Delta A^*$ | % error | $\langle W^*_{\text{dis}} \rangle$ |
| 0.001 | $2.1151 \pm 0.0002$ | 0.0006 | $0.0017 \pm 0.0003$ |
| 0.005 | $2.1149 \pm 0.0004$ | $-0.008$ | $0.0084 \pm 0.0006$ |
| 0.01 | $2.1147 \pm 0.0006$ | $-0.02$ | $0.0169 \pm 0.008$ |
| 0.02 | $2.1144 \pm 0.0008$ | $-0.03$ | $0.033 \pm 0.001$ |
| 0.05 | $2.116 \pm 0.001$ | 0.04 | $0.081 \pm 0.002$ |
| 0.1 | $2.114 \pm 0.002$ | $-0.06$ | $0.155 \pm 0.003$ |
| 0.2 | $2.116 \pm 0.003$ | 0.05 | $0.281 \pm 0.004$ |
| 0.5 | $2.116 \pm 0.004$ | 0.04 | $0.508 \pm 0.005$ |
| 1.0 | $2.115 \pm 0.005$ | $-0.01$ | $0.673 \pm 0.006$ |
| | Parameter set 2 : | $\Delta A^*_{\text{num}} = 5.55479$ | |
| $v^*$ | $\Delta A^*$ | % error | $\langle W^*_{\text{dis}} \rangle$ |
| 0.001 | $5.5551 \pm 0.0002$ | 0.005 | $0.0031 \pm 0.0004$ |
| 0.005 | $5.5559 \pm 0.0006$ | 0.02 | $0.0156 \pm 0.0008$ |
| 0.01 | $5.5546 \pm 0.0008$ | $-0.003$ | $0.031 \pm 0.001$ |
| 0.02 | $5.553 \pm 0.001$ | $-0.02$ | $0.063 \pm 0.002$ |
| 0.05 | $5.554 \pm 0.002$ | $-0.01$ | $0.155 \pm 0.003$ |
| 0.1 | $5.551 \pm 0.003$ | $-0.07$ | $0.306 \pm 0.004$ |
| 0.2 | $5.549 \pm 0.005$ | $-0.08$ | $0.593 \pm 0.006$ |
| 0.5 | $5.54 \pm 0.01$ | $-0.26$ | $1.38 \pm 0.01$ |
| 1.0 | $5.50 \pm 0.03$ | $-0.95$ | $2.42 \pm 0.03$ |

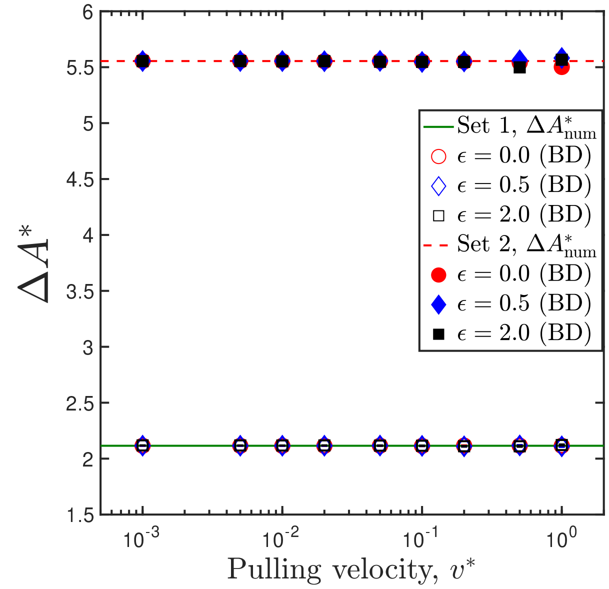


FIG. 1. Comparison of the numerically calculated free energy differences (indicated by horizontal lines) against that calculated using the JE, for two different parameter sets shown in Table I. Error bars represent a statistical uncertainty of one standard error of the mean (s.e.m).

### III. Parameter-Space specification

The molecular parameters in this study are the finite extensibility parameter, $b$, the length scale, $l_H$, the bead radius, $a$, and the damping coefficient of the dashpot, $K$. The control parameters are the stiffness of the traps, the initial extension of the molecule subjected to stretching, and the pulling velocity.

#### A. Molecular parameters

The choice of model parameters is based on the $\lambda$-phage DNA (48.5 kbp) used in Murayama et al.'s[8] work, which has a contour length, $L_0$, of $16.5\,\mu$m, and Kuhn segment length, $b_K$, of approximately 88 nm. In order to model this molecule as a dumbbell, the model parameters, $b$, and $l_H$, are chosen such that the contour length and the radius of gyration of the model and the DNA molecule are the same. Firstly, the number of Kuhn steps, $N_K$, in the DNA molecule is found using the relation,

$$L_0 = N_K b_K \tag{46}$$

to be $N_K \approx 188$. This quantity can be used to estimate the radius of gyration at theta conditions, $R_g^\theta$, as

$$R_g^\theta = \frac{\sqrt{N_K} b_K}{\sqrt{6}} = 492.6\,\text{nm} \tag{47}$$

The maximum permissible length of the dumbbell is

$$Q_0 \equiv L_0 = (N_b - 1)\sqrt{b} l_H = \sqrt{b}\, l_H \tag{48}$$

and the radius of gyration at theta conditions for the dumbbell is given by

$$\left(R_{\rm g}^{\theta}\right)^2 = \Gamma^2(b)\,\frac{N_{\rm b}^2-1}{2N_{\rm b}}\,l_{\rm H}^2 = \frac{3}{4}\Gamma^2(b)\,l_{\rm H}^2 \tag{49}$$

where $\Gamma(b)$ is a known function of $b$ for a given spring force law, and has been discussed extensively in ref. 9. Squaring Eq. (48) and dividing by Eq. (49), one can write

$$\Upsilon \equiv \frac{b}{3\Gamma^2(b)} = \frac{1}{4}\frac{\left(L_0\right)^2}{\left(R_{\rm g}^{\theta}\right)^2} = \frac{3}{2}N_{\rm K} \tag{50}$$

where the experimentally obtained quantities have been substituted in place of the model values of $L_0$ and $R_{\rm g}^{\theta}$. In ref. 9, the following expression has been obtained for a spring obeying the WLC force law,

$$b = 3\Upsilon\left[1 - \frac{0.63}{\sqrt{\Upsilon}} - \frac{1.0}{\Upsilon} - \frac{0.70}{\Upsilon^2} + \frac{1.47}{\Upsilon^3}\right] \tag{51}$$

Using Eq. (50) and Eq. (51), one obtains a value for $b$. Knowing the value of $b$ and $\Upsilon$, $\Gamma$ can be computed. Inserting the value for $\Gamma$ into Eq. (49), $l_{\rm H}$ can also be determined. Following this procedure for $\lambda$-phage DNA, we obtain $b = 811.25$ and $l_{\rm H} = 580.95\,$nm. We round down both these values, and use $b = 800$ and $l_{\rm H} = 500\,$nm as the parameters to model $\lambda$-phage DNA. The Hookean spring constant of the model, $H$, is then found using

$$H = \frac{k_BT}{l_{\rm H}^2} = \frac{4.142\ \text{pN nm}}{(500)^2\,(\text{nm})^2} = 1.657\times10^{-5}\,\text{pN/nm}$$

The choice of the bead radius, $a$, is motivated by Alexander-Katz et al.'s[10] work, where it is suggested that the monomeric radius may be taken as the persistence length of the molecule. For the DNA molecule considered here, the persistence length would be $L_{\rm p} \approx b_{\rm K}/2 = 44$ nm. We choose $a = 30\,$nm, which is close to this value and identical to the choice made by Alexander-Katz et al.[10] for comparing the results of BD simulations against experiments on DNA.

Other values of $b$ and $l_{\rm H}$ of the same order-of-magnitude as obtained for the $\lambda$-phage DNA case have also been used in the present study. In addition to $a = 30\,$nm, bead radii of 80 nm and 100 nm have also been used in this study.

Murayama et al.[8] estimate the internal friction coefficient of the DNA molecule in their experiments to have a value $K \approx 10^{-7}$ kg/s. Since their calculation of the drag coefficient using slender-body hydrodynamics proved to be too small to account for the dissipation observed in the stretch-relaxation process, they attribute the entire dissipation to the internal friction within the molecule. A choice of $K = 1.0\times10^{-7}\,$kg/s leads to the internal friction parameter having a value $\epsilon = 88$, necessitating the use of a longer equilibration time than that used in our simulations of $t^*_{\rm eqb} = 55.0$. For the purposes of the present study, values for $K$ between $1.0\times10^{-9}$ kg/s and $1.0\times10^{-8}$ kg/s have been chosen.

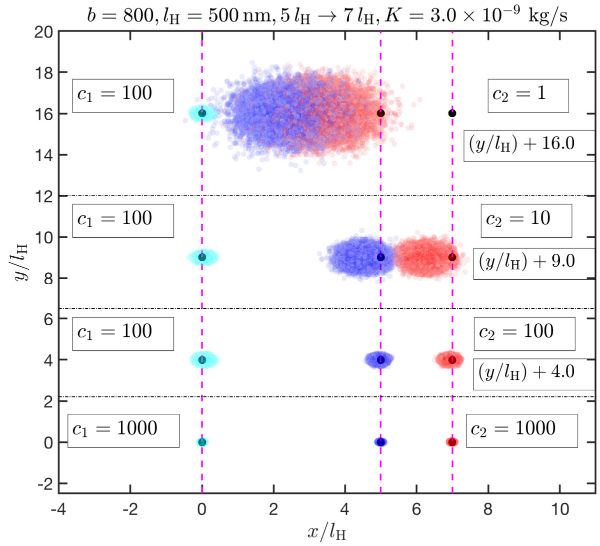


FIG. 2. An x-y projection of the equilibrated positions of the beads, for an ensemble of $N = 1\times10^4$ dumbbells, for different values of the trap stiffness. Data points in cyan correspond to the positions of the first bead with the corresponding trap at the origin, those in blue denote the positions of the second bead when the trap position is at $\boldsymbol{\chi}_2^{(\rm i)} = (5\,l_{\rm H}, 0, 0)$, and those in red represent the positions of the second bead when the trap is located at $\boldsymbol{\chi}_2^{(\rm f)} = (7\,l_{\rm H}, 0, 0)$. The dashed vertical lines, from left to right, represent the position of the first trap, the initial position of the second trap, and the final position of the second trap, respectively. The data sets have been shifted vertically for clarity, and the offsets for each of the shifted cases are indicated in the figure.

### B. Trap stiffness

Fig. 2 provides a snapshot of the $x$-$y$ projection of the positions of the beads of the dumbbell, obtained after an equilibration of fifty-five dimensionless times at the initial trap positions $\boldsymbol{\chi}_1^* = (0,0,0)\,,\ \boldsymbol{\chi}_2^{(\rm i)*} = (5,0,0)$ and final trap states $\boldsymbol{\chi}_1^* = (0,0,0)\,,\ \boldsymbol{\chi}_2^{(\rm f)*} = (7,0,0)$, as a function of the optical trap stiffness.

It is clearly seen that the strength of the trap ($c_1$ or $c_2$) determines its ability to confine the bead near the position of its minimum. A consequence of using "soft" traps is that the position of the beads in the trap undergo large fluctuations, and the distance over which the dumbbell is stretched does not correspond to the distance over which the trap is moved.

The dissipated work in the extrapolated limit of zero solvent viscosity is defined as the restoring force provided by the dashpot, multiplied by the distance over which the trap is moved (Eq. (1) of main text). When softer traps are used, not all of the trajectories correspond to an increase in the extension of the dumbbell, even though the average extension increases. The pulling efficiency may be defined as the fraction of the total trajectories that correspond to an increase in the extension of the spring from the initial stretch ($Q^{(\rm f)}$) to the final stretch

TABLE III. Effect of trap stiffness on pulling efficiency. The representative case considered here corresponds to stretching a molecule with parameters: $\{b = 800, l_{\mathrm{H}} = 500\,\mathrm{nm}, K = 3\times10^{-9}\,\mathrm{kg/s}\}$ from an initial trap position of $\boldsymbol{\chi}_2^{(\mathrm{i})} = (5\,l_{\mathrm{H}}, 0, 0)$ to a final trap position of $\boldsymbol{\chi}_2^{(\mathrm{f})} = (7\,l_{\mathrm{H}}, 0, 0)$. Results are obtained from simulations of $N = 1\times10^4$ trajectories, except when mentioned otherwise.

| Trap stiffness | $\left[\sqrt{\langle Q^2\rangle}\right]^{(\mathrm{f})*} - \left[\sqrt{\langle Q^2\rangle}\right]^{(\mathrm{i})*}$ | Pulling efficiency[b] (%) |
|---|---|---|
| $c_1 = 100,\, c_2 = 1$[a] | $0.843 \pm 0.003$ | 81.978 |
| $c_1 = 100,\, c_2 = 10$[a] | $1.739 \pm 0.001$ | 99.995 |
| $c_1 = c_2 = 100$[a] | $1.9452 \pm 0.0006$ | 100.0 |
| $c_1 = c_2 = 1000$ | $1.9938 \pm 0.0006$ | 100.0 |
| $c_1 = c_2 = 10000$ | $1.9993 \pm 0.0002$ | 100.0 |

[a] $N = 1\times10^5$
[b] Eq. (52)

($Q^{(\mathrm{i})}$), when the mobile trap is moved from its initial location to the final location :

$$\text{Pulling efficiency}(\%) = 100 \times \frac{\text{No. of trajectories with } Q^{(\mathrm{f})} > Q^{(\mathrm{i})}}{\text{Total number of trajectories}} \tag{52}$$

where $Q = |\boldsymbol{Q}|$.

In Table III, the effect of trap stiffness on the pulling efficiency for a representative case is presented. From the table, we note that when a trap stiffness of $c_1 = 100$ and $c_2 = 1$ is used, only $\sim 82\%$ of the total number of trajectories correspond to an increase in the extension of the dumbbell. In other words, there is no pulling work done against the spring-dashpot system for the remaining $\sim 18\%$ of the trajectories. The extraction of the internal friction coefficient from the average work calculated in this case produces erroneous results.

On the other hand, as the stiffness of the trap is increased, say to $c_1 = c_2 = 100$, *all* of the trajectories correspond to an increase in extension of the dumbbell, even though the increase in the extension of the dumbbell is not exactly the same as the distance over which the trap is moved. Nonetheless, the internal friction coefficient calculated in this case produces more accurate results in comparison to the soft pulling case, as discussed further in the context of Fig. 4 below.

In Fig. 3, for representative values of the molecular and control parameters, the first trap strength in dimensionless units is kept fixed, at $c_1 = 1000$, and the strength of the second trap is varied. As seen from the figure, the dissipation increases with an increase in the strength of the second trap, and reaches its saturation value at $c_2 \approx 100$. Since it is desirable to operate in a regime where the dissipated work is independent of the trap strength, we choose $c_1 = c_2 = 1000$ for all our simulations, except when specified otherwise.

In Fig. 4, the average dissipated work in the extrapolated limit of $\eta_s \to 0$, divided by the distance over which the trap is moved, is plotted against the pulling velocity, for a representative case, for various values of the trap stiffness. The slope of the graph gives the internal friction coefficient. As seen from the figure, the slopes of the curves increase with an increase in trap stiffness, and saturate once $c_1 = c_2 = 1000$ has been reached.

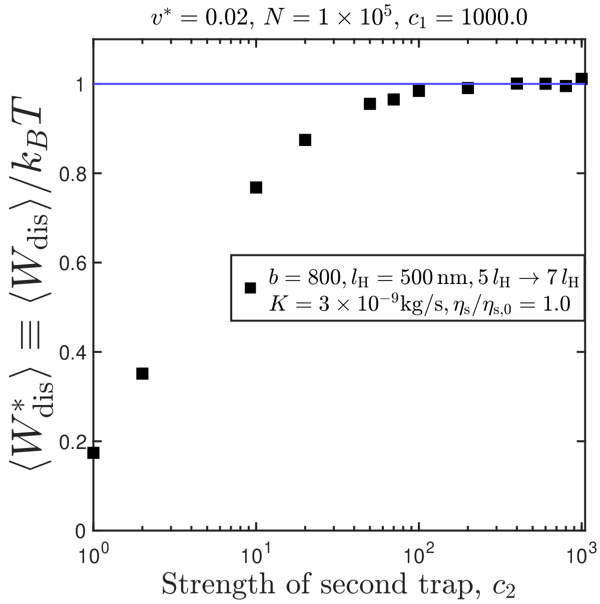


FIG. 3. Average dissipated work at a fixed value of the first trap stiffness, $c_1 = 1000.0$, as a function of pulling velocity, for various values of the second trap stiffness, $c_2$. Error bars represent a statistical uncertainty of one standard error of the mean (s.e.m).

The exact values of the coefficients for the various trap stiffnesses have been listed in Table IV. Also presented in the table are the extracted internal friction coefficients, when the definition of $d$ is changed to denote the change in the mean extension of the spring, i.e., $d = \left[\sqrt{\langle Q^2\rangle}\right]^{(\mathrm{f})} - \left[\sqrt{\langle Q^2\rangle}\right]^{(\mathrm{i})}$. Such a change in definition marginally improves the accuracy of the analysis for the soft trap cases, but has no effect for the stiff pulling cases because the two definitions for $d$ are nearly equivalent in

TABLE IV. Effect of trap stiffness on the estimation of internal friction coefficients. The representative case considered here corresponds to stretching a molecule with $\{b = 800, l_{\rm H} = 500\,{\rm nm}\}$ from an initial trap position of $\boldsymbol{\chi}_2^{({\rm i})} = (5\,l_{\rm H}, 0, 0)$ to a final trap position of $\boldsymbol{\chi}_2^{({\rm f})} = (7\,l_{\rm H}, 0, 0)$. Results are obtained from simulations of $N = 1 \times 10^4$ trajectories, except when mentioned otherwise. The error associated with the protocol is calculated as, % error $= 100 \times [(K_{\rm BD} - K)/K]$.

| | $d = \chi_{2x}^{({\rm f})} - \chi_{2x}^{({\rm i})} = 2\,l_{\rm H}$ | | |
|---|---|---|---|
| Trap stiffness | Input, $K[\times 10^9$ kg/s] | $K_{\rm BD}[\times 10^9$ kg/s] | % error |
| $c_1 = 100,\ c_2 = 1$[a] | 3.0 | $0.5730 \pm 0.0007$ | -80.90 |
| $c_1 = 100,\ c_2 = 10$[a] | 3.0 | $2.291 \pm 0.008$ | -23.63 |
| $c_1 = c_2 = 100$[a] | 3.0 | $2.858 \pm 0.008$ | -4.73 |
| $c_1 = c_2 = 1000$ | 3.0 | $2.95 \pm 0.02$ | -1.74 |
| $c_1 = c_2 = 10000$ | 3.0 | $2.96 \pm 0.01$ | -1.44 |
| $c_1 = 100,\ c_2 = 1$[a] | 6.0 | $1.124 \pm 0.003$ | -81.26 |
| $c_1 = 100,\ c_2 = 10$[a] | 6.0 | $4.579 \pm 0.006$ | -23.68 |
| $c_1 = c_2 = 100$[a] | 6.0 | $5.699 \pm 0.003$ | -5.01 |
| $c_1 = c_2 = 1000$ | 6.0 | $6.00 \pm 0.07$ | 0.11 |
| $c_1 = c_2 = 10000$ | 6.0 | $6.11 \pm 0.04$ | 1.83 |
| | $d = \left[\sqrt{\langle Q^2 \rangle}\right]^{({\rm f})} - \left[\sqrt{\langle Q^2 \rangle}\right]^{({\rm i})}$ | | |
| Trap stiffness | Input, $K[\times 10^9$ kg/s] | $K_{\rm BD}[\times 10^9$ kg/s] | % error |
| $c_1 = 100,\ c_2 = 1$[a] | 3.0 | $1.360 \pm 0.002$ | -54.65 |
| $c_1 = 100,\ c_2 = 10$[a] | 3.0 | $2.635 \pm 0.009$ | -12.15 |
| $c_1 = c_2 = 100$[a] | 3.0 | $2.927 \pm 0.01$ | -2.41 |
| $c_1 = c_2 = 1000$ | 3.0 | $2.96 \pm 0.02$ | -1.43 |
| $c_1 = c_2 = 10000$ | 3.0 | $2.96 \pm 0.01$ | -1.41 |
| $c_1 = 100,\ c_2 = 1$[a] | 6.0 | $2.893 \pm 0.008$ | -51.78 |
| $c_1 = 100,\ c_2 = 10$[a] | 6.0 | $5.268 \pm 0.007$ | -12.19 |
| $c_1 = c_2 = 100$[a] | 6.0 | $5.858 \pm 0.003$ | -2.37 |
| $c_1 = c_2 = 1000$ | 6.0 | $6.02 \pm 0.07$ | 0.38 |
| $c_1 = c_2 = 10000$ | 6.0 | $6.11 \pm 0.04$ | 1.89 |

[a] $N = 1 \times 10^5$

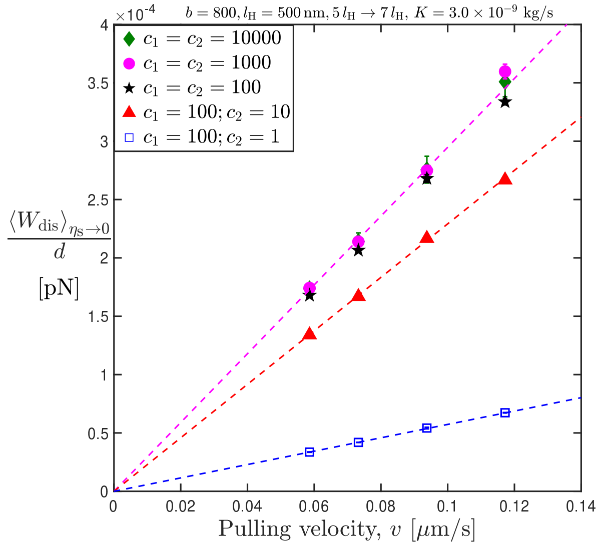


FIG. 4. Average dissipated work in the limit of zero solvent viscosity, divided by the stretching distance $d = \chi_2^{({\rm f})} - \chi_2^{({\rm i})}$, as a function of pulling velocity, for various values of the trap stiffness. Error bars represent a statistical uncertainty of one standard error of the mean (s.e.m).

the latter case.

#### C. Initial stretch of molecule

The accuracy of the protocol for extracting internal friction is clear from Table V. For all experiments in the linear regime of the force-extension profile, trap stiffnesses of $c_1 = c_2 = 1000$ suffice to yield a value for the internal friction coefficient that is within 2% of the actual value. Parameter set (A) appeared to be an exception, but yielded accurate results when simulations were performed over a larger ensemble size.

On the other hand, for starting positions that are in the non-linear regime of the force-extension profile, the default trap stiffness of $c_1 = c_2 = 1000$ yields results with greater than 5% error. Increasing the ensemble size leads to more precise results, but does not improve the accuracy. A ten-fold increase in the trap stiffnesses, however, results in an improved accuracy in the predictions, bringing the error lower than 5%.

In Fig. 5, the force-extension profile of the Marko-Siggia expression is plotted. The force-law is linear at low values of the fractional extension, and diverges as the fractional extension approaches unity. The various

TABLE V. Internal friction coefficients estimated from our protocol, for various values of the molecular parameters, and initial extension of the molecule. Trap stiffnesses of $c_1 = c_2 = 1000$, and an ensemble size of $N = 1 \times 10^4$ trajectories was used for all entries, except when specified otherwise. The error associated with the protocol is calculated as, % error $= 100 \times [(K_{\rm BD} - K)/K]$.

| | Parameters | Input, $K[\times 10^9$ kg/s] | $K_{\rm BD}[\times 10^9$ kg/s] | % error |
|---|---|---|---|---|
| (A) | $b = 200$, $l_{\rm H} = 150\,$nm, $d = l_{\rm H}$ | 1.0 | $1.04 \pm 0.01$ | 4.42 |
| (A)[a] | $b = 200$, $l_{\rm H} = 150\,$nm, $d = l_{\rm H}$ | 1.0 | $0.996 \pm 0.004$ | −0.40 |
| (B) | $b = 200$, $l_{\rm H} = 150\,$nm, $d = l_{\rm H}$ | 10.0 | $10.04 \pm 0.05$ | 0.47 |
| (C) | $b = 800$, $l_{\rm H} = 500\,$nm, $d = 2\,l_{\rm H}$ | 3.0 | $2.95 \pm 0.02$ | −1.74 |
| (D) | $b = 800$, $l_{\rm H} = 500\,$nm, $d = 2\,l_{\rm H}$ | 6.0 | $6.00 \pm 0.07$ | 0.11 |
| (E) | $b = 400$, $l_{\rm H} = 350\,$nm, $d = 2\,l_{\rm H}$ | 1.0 | $0.99 \pm 0.03$ | −1.36 |
| (F) | $b = 400$, $l_{\rm H} = 350\,$nm, $d = 2\,l_{\rm H}$ | 10.0 | $9.922 \pm 0.008$ | −0.78 |
| (G) | $b = 200$, $l_{\rm H} = 150\,$nm, $d = l_{\rm H}$ | 1.0 | $0.942 \pm 0.005$ | −5.83 |
| (G)[a] | $b = 200$, $l_{\rm H} = 150\,$nm, $d = l_{\rm H}$ | 1.0 | $0.935 \pm 0.003$ | −6.48 |
| (G)[b] | $b = 200$, $l_{\rm H} = 150\,$nm, $d = l_{\rm H}$ | 1.0 | $0.97 \pm 0.01$ | −3.22 |
| (H) | $b = 200$, $l_{\rm H} = 150\,$nm, $d = l_{\rm H}$ | 10.0 | $9.4 \pm 0.1$ | −6.31 |
| (H)[a] | $b = 200$, $l_{\rm H} = 150\,$nm, $d = l_{\rm H}$ | 10.0 | $9.224 \pm 0.006$ | −7.76 |
| (H)[b] | $b = 200$, $l_{\rm H} = 150\,$nm, $d = l_{\rm H}$ | 10.0 | $9.85 \pm 0.06$ | −1.49 |

[a] $N = 1 \times 10^5$
[b] $c_1 = c_2 = 10000$

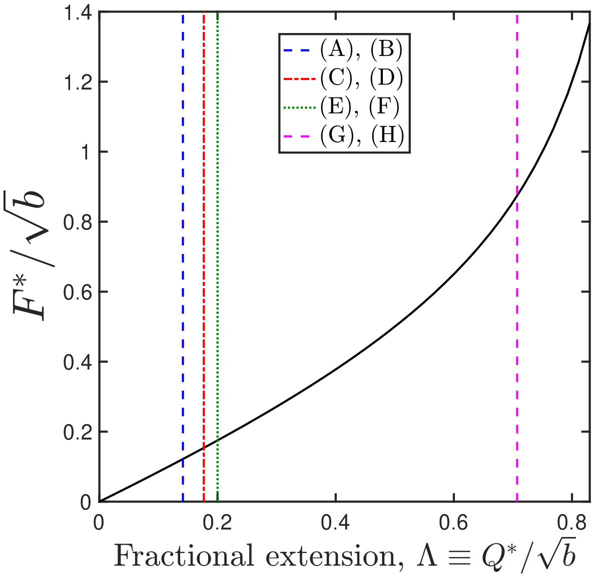


FIG. 5. Force extension profile for the Marko-Siggia force expression. Vertical lines indicate the fractional initial stretch, $\left\{\chi_2^{(\rm i)*}/\sqrt{b}\right\}$, of the molecule that is subject to pulling. Starting from the fractional extension indicated by the vertical lines, the molecule is pulled over a distance $d$. The parameters for the various molecules subjected to pulling, and their respective pulling distances, are provided in Table V.

pulling experiments performed in this work are symbolically indicated by means of vertical lines in the plot. This figure must be read in conjunction with Table V wherein the molecular parameters, and the stretching distances pertaining to each vertical line are given. For example, the green dotted vertical line corresponds to the parameter set (E). This means that a molecule straddled between the two traps, such that the dimensionless distance between the two traps normalised by the dimensionless contour length is initially 0.2, and is pulled over a distance $d = 2\,l_{\rm H}$.

#### D. Pulling velocity and ensemble size

The Jarzynski equality (JE), $\langle \exp[-W^*] \rangle = \exp[-\Delta A^*]$, is strictly *exact* only in the limit of an infinite number of work trajectories, $N \to \infty$. In applications of the JE, the number of trajectories required to accurately recover the free-energy difference increases with average dissipated work $[\langle W^*_{\rm dis} \rangle = \langle W^* \rangle - \Delta A^*]$ in the process, as discussed in refs. 11–13.

In Figs. 6, the effect of internal friction and pulling velocity on the probability distribution of the work trajectories is plotted. The vertical green lines in the figures indicate the free-energy difference, $\Delta A^*$, obtained by taking an error-weighted mean of the values of the free energy difference obtained at dimensionless pulling velocities $(v^*) \leq 0.02$.

From Fig. 6 (a), it is seen that increasing the pulling velocity at a fixed value of the internal friction parameter increases the average dissipated work, and the width of the distribution. An identical trend is observed in Fig. 6 (b), where an increase in the internal friction parameter at a fixed pulling velocity causes the work distribution to shift rightwards, and results in an increased dissipation. Thus, the dissipation in our model is directly correlated with the pulling velocity, and the internal friction in the system.

Under such conditions of high dissipation, the estimates for $\Delta A$ are dominated by rare realizations that occur near the tail of the work distribution, necessitating

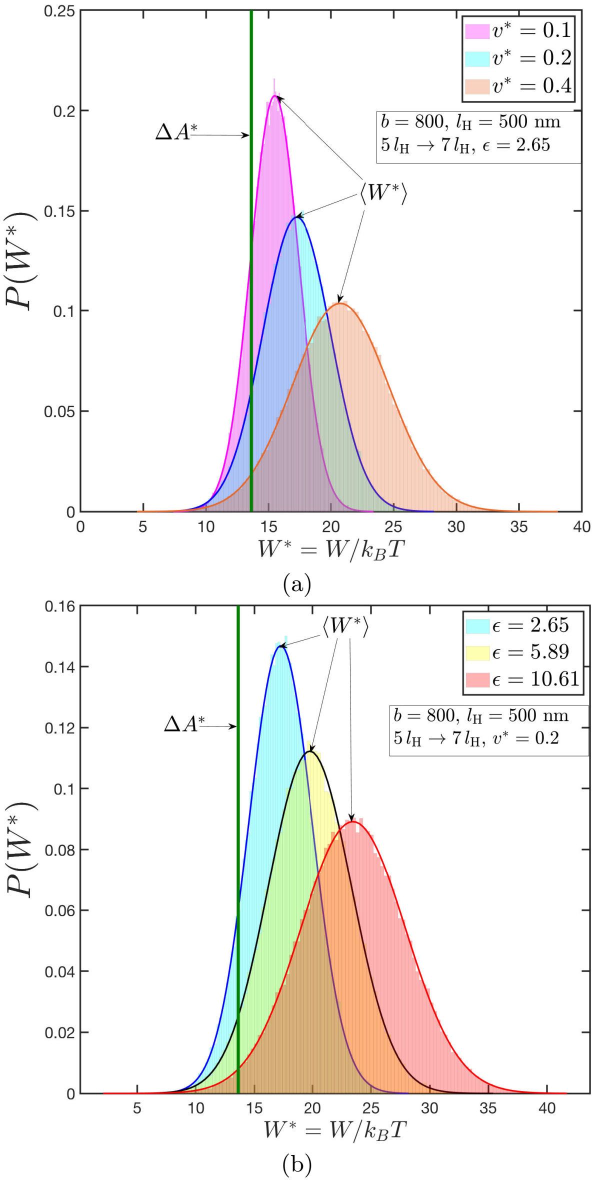


FIG. 6. Probability densities of the work done over $10^5$ realizations of the pulling protocol, for: (a) a fixed value of the internal friction parameter, and three different values of the pulling velocity, and (b) a fixed pulling velocity, and three different values of the internal friction parameter. The green vertical line represents the free-energy difference obtained by taking an error-weighted mean of the values of the free energy difference obtained at pulling velocities $(v^*) \leq 0.02$ using JE.

the use of a larger number of trajectories to obtain an accurate estimate of the free energy difference.

Without prior knowledge of the ensemble size required for the simulations, an initial guess of $N = 1 \times 10^4$ was chosen. The plot of dissipated work against the pulling velocity for this choice of ensemble size is presented in Fig. 7. The averaged dissipated work scales nearly linearly with the pulling velocity for all the cases, except for the data set with the highest internal friction parameter.

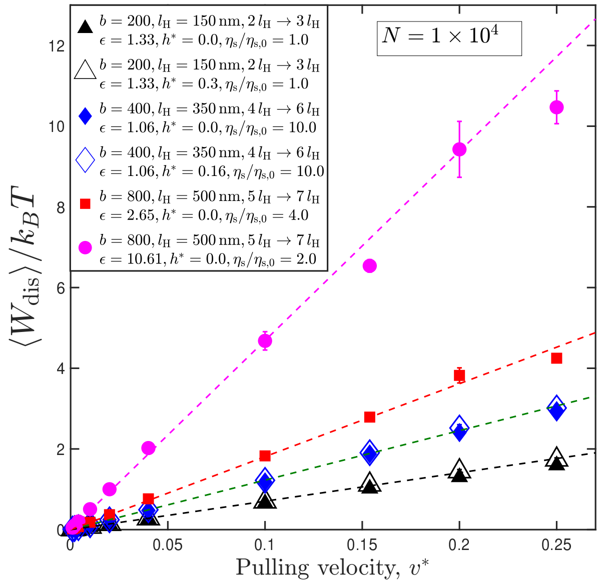


FIG. 7. Average dissipated work, as a function of pulling velocity, for various values of the molecular and control parameters and an ensemble size of $N = 1 \times 10^4$. Trap stiffness used is $c_1 = c_2 = 1000$. Symbols indicating datasets with fluctuating hydrodynamic interactions have been enlarged for clarity. Error bars, which represent a statistical uncertainty of one standard error of the mean (s.e.m), are smaller than the symbol size.

Upon closer inspection of that dataset, as presented in Fig. 8, it is found that the deviation from linearity is due to the incorrect estimation of the free energy difference at higher pulling velocities, when an ensemble size of $N = 1 \times 10^4$ is used. Upon increasing the ensemble size for the higher velocity cases *empirically*, the accuracy of the estimated free energy difference improves. The ensemble size is plotted as a function of the average dissipated work in the inset of Fig. 8.

Therefore, it is seen that the choice of the pulling velocity and the ensemble size are mutually related.

#### E. Experimental bounds on optical tweezer parameters

In Table VI, based on a survey of the literature, the range of trap stiffnesses, pulling velocities, and stretching distances typically accessible by optical tweezers is given. Additionally, it is instructive to examine the typical values of dissipation and ensemble sizes encountered in optical tweezer induced unfolding experiments. Liphardt et al.[14] stretch RNA hairpins using optical tweezers, and estimate $\langle W_{\mathrm{dis}} \rangle = 2\text{-}3\,k_BT$ with $N = 47$. Similarly, for pulling experiments on DNA hairpins performed by Gupta et al.,[15] $\langle W_{\mathrm{dis}} \rangle = 1.1 \pm 0.7 k_BT$ for $N = 99$, and $\langle W_{\mathrm{dis}} \rangle = 4.9 \pm 0.3 k_BT$ for $N = 1293$. As discussed previously, higher values of dissipation necessitate the use of larger ensemble sizes.

Furthermore, the position and force resolution limits

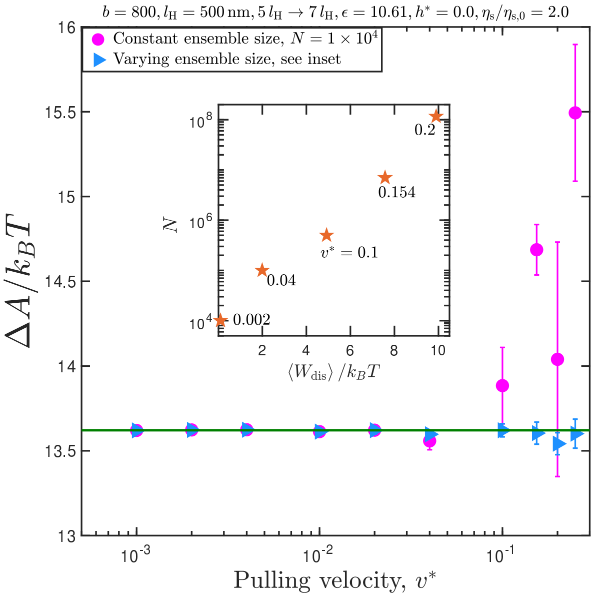


FIG. 8. Free energy difference as a function of pulling velocity, for a representative case. Inset shows the empirically chosen ensemble size as a function of the average dissipated work. The numbers next to the data points indicate the corresponding values of the dimensionless pulling velocity. The green horizontal line represents the free-energy difference obtained by taking an error-weighted mean of the values of the free energy difference obtained at pulling velocities $(v^*) \leq 0.02$ using Jarzynski's equality. Trap stiffness used is $c_1 = c_2 = 1000$. Error bars represent a statistical uncertainty of one standard error of the mean (s.e.m).

of optical traps are inversely correlated: stiffer traps improve the spatial resolution but also introduce large fluctuations in the measured force. A rough estimate of these resolution limits may be obtained using the equipartition theorem, as explained in refs. 18,19. Most commercial optical tweezer setups are equipped with filtering mechanisms that aid in improving the precision in the measurements, by reducing the resolution limits.[15] A detailed discussion of the resolution offered by optical tweezers can be found in refs. 18 and 19.

TABLE VI. Typically observed lower and upper bounds on optical tweezer parameters.

| Parameter | Lower bound | Upper bound |
|---|---|---|
| Trap stiffness (pN/nm) | 0.0002 [ref. 16] | 0.9 [ref. 15] |
| Pulling velocity (nm/s) | 10 [ref. 15] | 13560 [ref. 17] |
| Stretching distance (nm) | 10 [ref. 15] | 8000 [ref. 8] |

### IV. Error bars

Let $y_i$ represent a quantity that is obtained from the $i$th trajectory of a Brownian dynamics simulation. The ensemble average of $y$ over $N$ trajectories is then calculated as,

$$\langle y \rangle = \frac{1}{N} \sum_{i=1}^{N} y_i \tag{53}$$

The statistical uncertainty in $\langle y \rangle$ is then quantified using the standard error of mean (s.e.m), calculated as[5]

$$\delta y = \sqrt{\frac{\left(\frac{\sum_i y_i^2}{N}\right) - \left(\frac{\sum_i y_i}{N}\right)^2}{(N-1)}} \tag{54}$$

where the summations run from $i = 1$ to $N$.

The statistical error bar associated with the free energy difference is evaluated as follows. In the dimensionless form, Jarzynski's equality is written as

$$\langle \exp\left[-W^*\right] \rangle = \exp\left[-\Delta A^*\right] \tag{55}$$

Therefore,

$$\Delta A^* = -\ln J; \quad J \equiv \langle \exp[-W^*] \rangle \tag{56}$$

$$\delta\left(\Delta F\right) = \left| -\frac{1}{J} \delta J \right| \tag{57}$$

where $\delta J$ is the s.e.m of $\langle \exp[-W^*] \rangle$, and $\delta\left(\Delta A^*\right)$ is the statistical error bar in $\Delta A^*$.

A linear least-squares fitting procedure is used for all the straight-line fits discussed in this work, and the statistical errors in the fitting parameters are obtained by taking a square root of the diagonal elements of the covariance matrix. The `fitlm` functionality of MATLAB is used for all the fitting exercises performed in the paper.


[1] R. B. Bird, C. F. Curtiss, R. C. Armstrong, and O. Hassager, *Dynamics of Polymeric Liquids - Volume 2 : Kinetic Theory* (John Wiley and Sons, 1987).
[2] R. Kailasham, R. Chakrabarti, and J. R. Prakash, J. Chem. Phys. **149**, 094903 (2018).
[3] J. Rotne and S. Prager, J. Chem. Phys. **50**, 4831 (1969).
[4] H. Yamakawa, *Modern Theory of Polymer Solutions* (Harper and Row, 1971).
[5] H. C. Öttinger, *Stochastic Processes in Polymeric Fluids* (Springer, 1996).
[6] W. Press, S. Teukolsky, W. Vetterling, and B. Flannery, *Numerical Recipes 3rd Edition: The Art of Scientific Computing* (Cambridge University Press, 2007).
[7] C. Jarzynski, Phys. Rev. Lett. **78**, 2690 (1997).
[8] Y. Murayama, H. Wada, and M. Sano, Eur. Phys. Lett. **79**, 58001 (2007).
[9] P. Sunthar and J. R. Prakash, Macromolecules **38**, 617 (2005).
[10] A. Alexander-Katz, H. Wada, and R. R. Netz, Phys. Rev. Lett. **103**, 028102 (2009).
[11] F. Ritort, C. Bustamante, and I. Tinoco, Proc. Natl. Acad. Sci. U.S.A. **99**, 13544 (2002).
[12] C. Jarzynski, Phys. Rev. E **73**, 046105 (2006).
[13] N. Yunger Halpern and C. Jarzynski, Phys. Rev. E **93**, 1 (2016).
